# Single-molecule live imaging of subunit interactions and exchange within cellular regulatory complexes

**DOI:** 10.1101/2024.06.25.600644

**Authors:** Thomas G.W. Graham, Claire Dugast-Darzacq, Gina M. Dailey, Britney Weng, Xavier Darzacq, Robert Tjian

**Affiliations:** Department of Molecular and Cell Biology and University of California, Berkeley; Howard Hughes Medical Institute, University of California, Berkeley; Université Paris Cité, CNRS, Institut Jacques Monod, F-75013 Paris, France

**Keywords:** proximity-assisted photoactivation (PAPA), fast single-molecule tracking (fSMT), transcriptional regulation, positive transcription elongation factor b (P-TEFb), protein-protein interactions, 7SK complex, heterogeneous nuclear ribonucleoproteins (hnRNPs)

## Abstract

Cells are built from vast networks of competing molecular interactions, most of which have been impossible to monitor in vivo. We recently devised a new strategy, proximity-assisted photoactivation (PAPA), to detect these interactions at single-molecule resolution in live cells. Here we apply PAPA to visualize the network of interactions that regulate the central transcription elongation factor P-TEFb. PAPA between multiple pairs of endogenous proteins, combined with fast single-molecule tracking (fSMT), revealed that inactive P-TEFb within the 7SK ribonucleoprotein complex is largely unbound to chromatin, that this complex dissociates within minutes of treatment with a P-TEFb kinase inhibitor, and that heterogeneous ribonucleoproteins (hnRNPs) bind 7SK concomitant with P-TEFb release. Unlike 7SK-bound P-TEFb, P-TEFb associated with the coactivator BRD4 exhibited increased binding to chromatin. Our results address longstanding questions about a key transcriptional regulator and demonstrate that PAPA-fSMT can probe subunit interactions and exchange within endogenous regulatory complexes in live cells.

## Introduction

Specific macromolecular interactions underlie all cellular regulatory mechanisms. A quintessential example is transcriptional regulation, which involves extensive networks of interactions between transcription factors, cofactors, RNA polymerase II (Pol II), and chromatin. One key transcriptional cofactor is the positive transcription elongation factor b (P-TEFb), a heterodimer of cyclin-dependent kinase 9 (Cdk9) and Cyclin T1/T2 that is broadly required to release Pol II from promoter-proximal pausing into productive elongation.^1–6^ P-TEFb is regulated by interactions with many other nuclear factors. Catalytically active P-TEFb is thought to be directed to target genes by interaction with BRD4,^7,8^ the super-elongation complex (SEC),^9^ and various sequence-specific transcription factors—both endogenous and viral.^10–17^ At the same time, a large fraction of P-TEFb is kept inactive within a nucleoprotein complex comprising the 7SK small nuclear RNA and the proteins HEXIM1/2, LARP7, and MEPCE.^18–24^ P-TEFb and its regulators have emerged as potential drug targets for treating HIV infection and various cancers.^25–30^

An important open question is how interaction of P-TEFb with these diverse partners is regulated. P-TEFb can be released from 7SK by activators such as HIV Tat and by various external stimuli, including DNA damage, inhibitors of Cdk9 and Pol II, and activation of signaling pathways that posttranslationally modify P-TEFb or HEXIM.^21,31–37^ Tat is thought to dissociate P-TEFb from 7SK by competitive interaction with both partners.^37–39^ Transcription inhibitors, including Cdk9 inhibitors, are proposed to displace P-TEFb from 7SK indirectly via promiscuous RNA-binding proteins called heterogeneous nuclear ribonucleoproteins (hnRNPs);^40–42^ this model proposes that blockage of nascent RNA synthesis causes accumulation of free hnRNPs, which bind 7SK and displace P-TEFb.^40,41^

In addition, some transcriptional activators have been proposed to bind the P-TEFb:7SK complex and stimulate local release of P-TEFb to activate nearby target genes.^43–47^ Supporting this model, ChIRP-seq of 7SK and ChIP-seq of HEXIM1 and LARP7 revealed widespread crosslinking across the genome,^43,48,49^ while RNA-FISH showed enrichment of 7SK at a synthetic reporter gene array,^50^ suggesting that P-TEFb:7SK is recruited to specific sites on chromatin. In contrast, cell lysis and fractionation experiments showed that the P-TEFb:7SK complex is soluble in a low-salt buffer, suggesting that it associates with chromatin weakly, if at all.^31,51^ Because these experiments all required either cell fixation or lysis, it remains unclear to what extent the P-TEFb:7SK complex associates with chromatin in vivo.

While P-TEFb interactions have been studied in cell lysates and crosslinked samples, it has so far been impossible to monitor them in real time in live cells. We recently described a new approach to detect protein interactions in live cells called proximity-assisted photoactivation (PAPA). In PAPA, excitation of a “sender” fluorophore reactivates a nearby “receiver” fluorophore from a photochemically-induced dark state (Figure 1 and Movie S1).^52–54^ This proximity-dependent reactivation can be used to detect interaction between a pair of proteins labeled with sender and receiver fluorophores. Unlike previous approaches, such as single-molecule FRET and two-color colocalization, PAPA has a flexible distance-dependence, is not impeded by spectral crosstalk, and can efficiently detect intracellular protein interactions at physiological concentrations in fast single-molecule tracking (fSMT), making it possible to localize and measure the mobility of complexes containing a specific pair of subunits in living cells.

**Figure 1:**
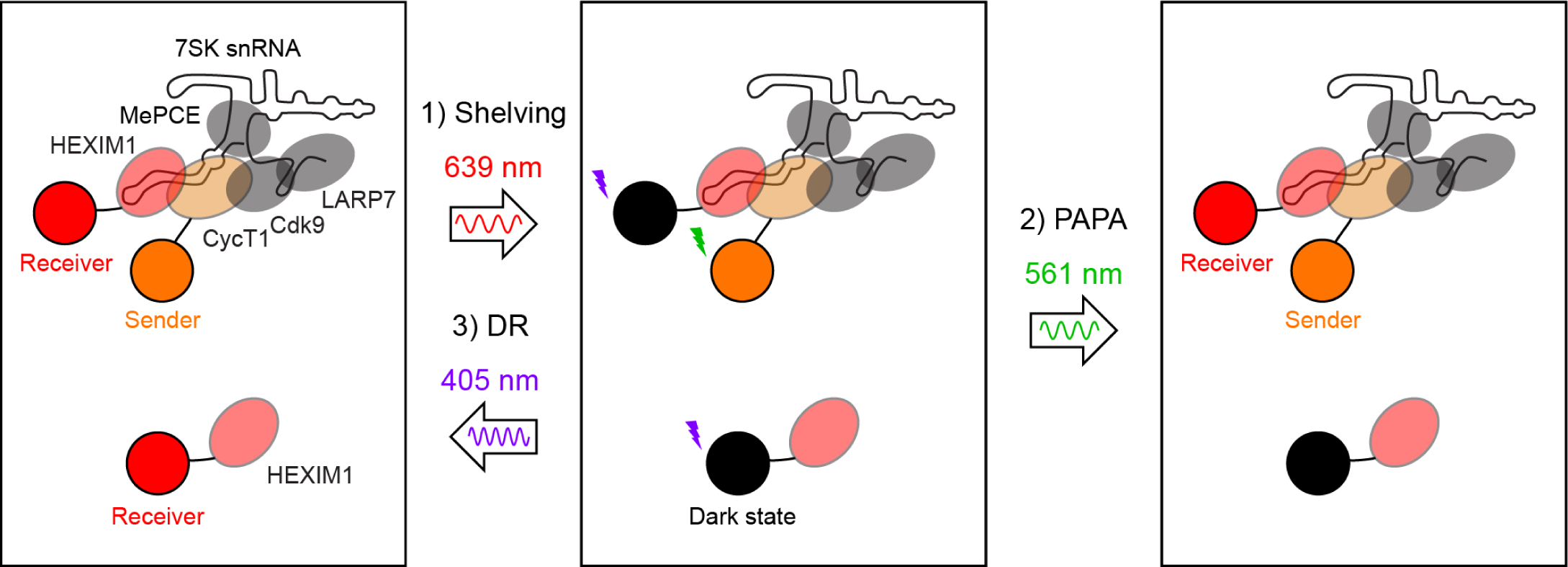
Detecting protein complexes with PAPA. A target protein fused to SNAPf (e.g., HEXIM1) is labeled with a receiver fluorophore (e.g., JFX650). A partner protein fused to Halo (e.g., CycT1) is labeled with a sender fluorophore (e.g., JFX549). *Step 1:* The receiver fluorophore is “shelved” in a photochemically-induced dark state by excitation with intense red (639 nm) light. About 10-15% of SNAPf-bound JFX650 fluorophores enter a dark state at this step, while 85-90% irreversibly photobleach.^55^ *Step 2:* Excitation of the sender with a brief pulse of green (561 nm) light selectively reactivates nearby receiver fluorophores by proximity-assisted photoactivation (PAPA). Reactivated receiver fluorophores are then imaged with red excitation. *Step 3:* As an internal control in the same cell, violet (405 nm) light is used to reactivate dark-state receiver fluorophores independent of their proximity to the sender. Multiple rounds of shelving, PAPA, and DR are performed for each cell, and the durations of green and violet pulses are adjusted for each protein pair to yield a density of reactivated molecules that is sufficiently sparse for tracking. Halo is labeled with the sender and SNAPf with the receiver, as labeling is more efficient for Halo, while dark state formation of the attached fluorophore is more efficient for SNAPf.^55^ Please note that this cartoon of 7SK is not meant to be structurally accurate, and for simplicity we have not shown that HEXIM1 forms a dimer.

Here, we applied PAPA and fSMT to probe the network of interactions that regulate P-TEFb. PAPA between four different pairs of sender- and receiver-labeled P-TEFb:7SK complex subunits revealed a select subpopulation of molecules with a characteristic diffusion coefficient. Interestingly, P-TEFb:7SK complexes were predominantly mobile, arguing against widespread stable tethering to chromatin. Time course PAPA measurements revealed rapid, synchronous dissociation of P-TEFb and HEXIM1 from the 7SK complex upon treatment with a P-TEFb inhibitor. PAPA detected interaction of multiple hnRNPs with the 7SK complex, and P-TEFb inhibitor treatment caused hnRNP R binding to 7SK to increase with kinetics similar to, if not slightly faster than P-TEFb:7SK dissociation. Finally, PAPA measurements detected dose-dependent dissociation of the P-TEFb:7SK complex by the HIV transcriptional activator Tat, while PAPA-fSMT showed that association of P-TEFb with the coactivator BRD4 is accompanied by increased interaction with chromatin.

These results reveal previously unmeasurable properties of P-TEFb regulatory complexes in their native context. More broadly, they demonstrate that PAPA-fSMT can detect endogenous protein complexes, measure their mobilities, and monitor their association and dissociation in live cells. PAPA-fSMT thus provides a useful means to probe dynamic protein interaction networks that regulate transcription and other cellular processes.

## Results

### Characterizing the endogenous P-TEFb:7SK complex using PAPA-fSMT

To visualize P-TEFb complexes in live U2OS cells, we fused endogenous CycT1 and HEXIM1 to the self-labeling Halo and SNAPf tags. Both Halo- and SNAPf-labeled CycT1 remained functional, based on induction of the HIV LTR promoter by the activator Tat, which is known to require CycT1 (Figure S1A).^56,57^ We tracked the motion of single molecules in live cells using stroboscopic HILO imaging and analyzed the resulting trajectories using Bayesian state array analysis, which infers the underlying distribution of molecular diffusion coefficients without presupposing a small number of discrete mobility states.^58,59^ Strikingly, for both CycT1 and HEXIM1, we observed three major subpopulations with distinct mobilities—an “immobile” fraction moving at ∼0.01 µm^2^/s (the approximate lower limit of our measurement, and the smallest diffusion coefficient in our state array), another at 2-3 µm^2^/s, and a third at 6-7 µm^2^/s (Figures 2A-B and S1B-C). We hypothesized that the latter two diffusing peaks correspond to distinct complexes of CycT1 and HEXIM1.

**Figure 2:**
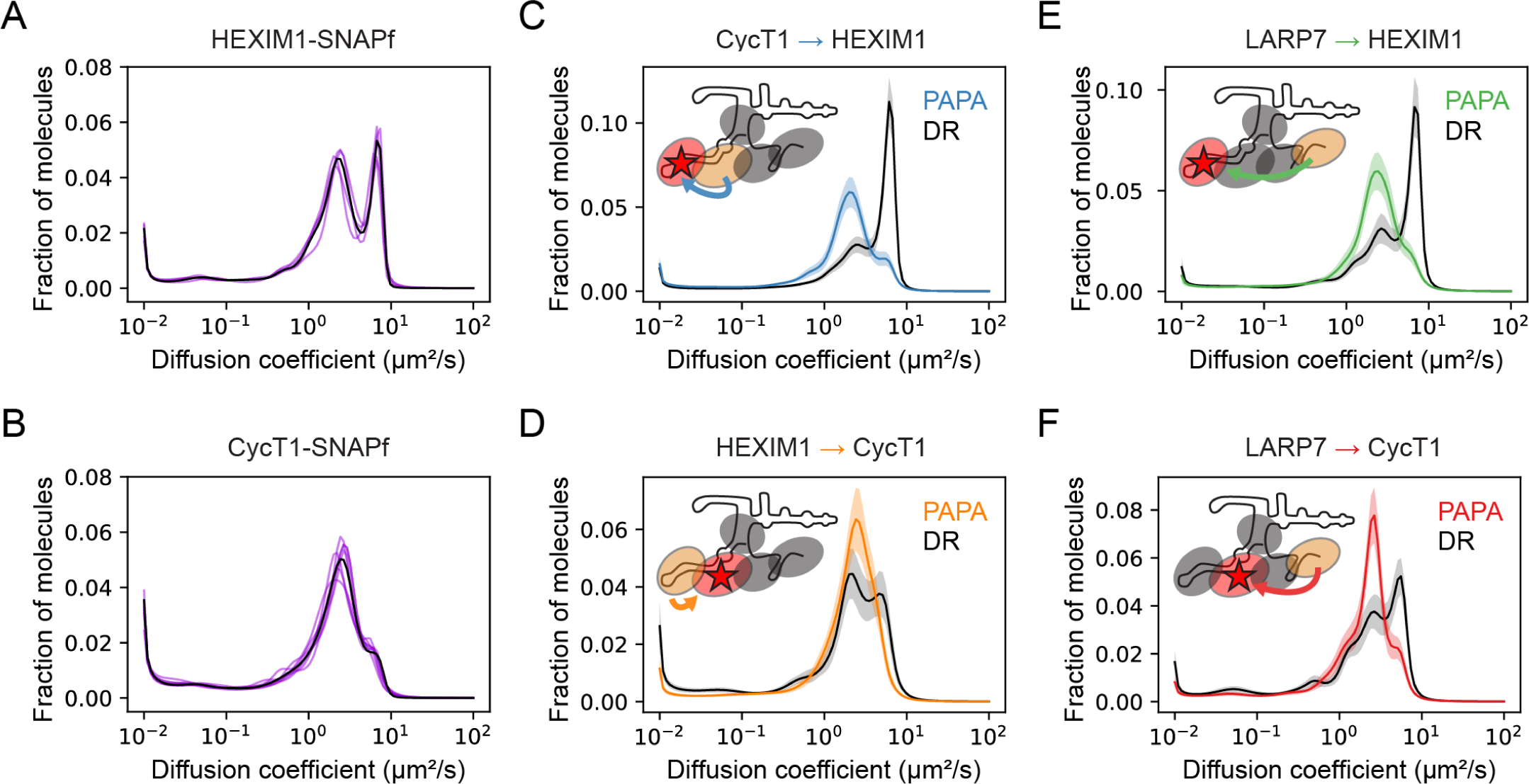
Visualization of the P-TEFb:7SK complex using PAPA-fSMT. **A-B)** Diffusion coefficient distributions from state array analysis of fSMT data for HEXIM1-SNAPf (panel A, clone S-A6) and SNAPf-CycT1 (panel B, clone G4-9). Individual experimental replicates are shown in transparent violet (n = 4 for HEXIM1-SNAPf and n = 7 for SNAPf-CycT1), while a combined analysis is shown in black. See also Figure S1B-C. **C-F)** Diffusion coefficient distributions of molecules reactivated by PAPA between the indicated sender → receiver pair (color curve) and direct reactivation (DR) of the receiver as an internal control (black curve). The cartoons (see Figure 1 for legend) represent which protein is labeled with the sender (orange) and which is labeled with the receiver (red). The red star emphasizes that the receiver-labeled protein is the one being tracked in single-molecule imaging. Solid curves and transparent regions represent the mean and approximate 95% confidence intervals (± 2 * S.D.) from bootstrapping analysis. **C)** Halo-CycT1 → HEXIM1-SNAPf (Clone HC-C6) **D)** HEXIM1-Halo → SNAPf-CycT1 (Clone SC-B12) **E)** LARP7-Halo → HEXIM1-SNAPf (Clone HHL-D6) **F)** LARP7-Halo → SNAPf-CycT1 (Clone CHL-D6).

To identify these complexes, we combined fSMT with proximity-assisted photoactivation (PAPA), in which a **sender** fluorophore on one protein is used to reactivate a **receiver** fluorophore on a second protein from a photochemically-induced dark state, making visible a specific complex of interest (Figure 1 and Movie S1).^52^ As a first step, we sought to visualize binding of the HEXIM1 subunit of the 7SK RNP complex to P-TEFb. To this end, we generated a double-homozygous Halo-CycT1 + HEXIM1-SNAPf endogenous knock-in U2OS cell line and labeled Halo-CycT1 with the PAPA sender fluorophore JFX549 and HEXIM1-SNAPf with the PAPA receiver fluorophore JFX650 (Figures 1, 2C, and S2A-F). After “shelving” the receiver in a dark state with intense red light (639 nm; Figure 1, Step 1), we applied short pulses of green light (561 nm) to excite sender-labeled CycT1 and reactivate nearby receiver-labeled HEXIM1 (Figure 1, Step 2). As an internal control, the same cell was exposed to pulses of violet light (405 nm) to induce direct reactivation (DR) of receiver-labeled HEXIM1, independent of proximity to CycT1 (Figure 1, Step 3). Each cell was imaged with several cycles of alternating PAPA, DR, and re-shelving/photobleaching (see Methods and Movie S1). Using this protocol, molecules reactivated by PAPA are enriched for complexes, while molecules reactivated by DR are somewhat depleted of complexes due to irreversible photobleaching of complexes reactivated by PAPA in preceding cycles.

Compared to HEXIM1 molecules reactivated by violet light (DR), HEXIM1 molecules reactivated by green light (PAPA) were enriched for the 2-3 µm^2^/s peak, implying that this subpopulation includes HEXIM1 bound to CycT1 (Figure 2C). Next, we performed the reciprocal HEXIM1 → CycT1 PAPA experiment using a double-homozygous HEXIM1-Halo + SNAPf-CycT1 cell line (Figure 2D). HEXIM1 → CycT1 PAPA likewise enriched a peak at 2-3 µm^2^/s, implying that this peak contains CycT1 bound to HEXIM1.

As an additional control, we quantified the PAPA signal using the “GV ratio”, defined as the ratio of the increase in the number of localizations induced by green light pulses (PAPA) to the increase induced by violet light pulses (DR). A background level of PAPA signal was observed between HEXIM1-Halo and two non-interacting control proteins, nuclear SNAPf (GV = 0.156 ± 0.007, 95% C.I.) and H2B-SNAPf (GV = 0.166 ± 0.012), which we attribute to chance proximity of sender- and receiver-labeled molecules.^55^ In contrast, a much higher GV ratio was observed between HEXIM1-Halo and SNAPf-CycT1 (3.36 ± 0.11), indicating specific interaction (Figure S3A).

*In vitro* biochemical results indicate that P-TEFb and HEXIM1 interact predominantly within the 7SK complex,^20,60,61^ and the much slower diffusion of PAPA-enriched HEXIM1 and CycT1 (Figure 2C-D) suggests incorporation in a large complex. To test further whether the 2-3 µm^2^/s subpopulation of CycT1 and HEXIM1 is associated with the 7SK complex, we next performed PAPA experiments between these two factors and the constitutive 7SK complex subunit LARP7. Fast SMT of Halo-tagged endogenous LARP7 revealed a major mobile population with a peak diffusion coefficient of 2.7 µm^2^/s (Figure S3C). Within LARP7-Halo cells, we then tagged CycT1 or HEXIM1 with SNAPf (Figure S2G-H) and used PAPA-fSMT to probe the interaction of each with LARP7. A higher PAPA signal was observed for SNAPf-tagged CycT1 and HEXIM1 than for a non-interacting H2B-SNAPf control, indicating interaction with LARP7 (Figure S3B). As predicted, both LARP7 → CycT1 and LARP7 → HEXIM1 PAPA enriched a subpopulation at 2-3 µm^2^/s (Figure 2C-D). Taken together, these PAPA-fSMT measurements show that endogenous P-TEFb, HEXIM1, and LARP7 interact within a discrete complex with a diffusion coefficient of 2-3 µm^2^/s.

Strikingly, only a small fraction of PAPA-reactivated P-TEFb:7SK complexes had the low mobility (D < 0.1 µm^2^/s) characteristic of chromatin binding (Figure 2C-F). This immobile fraction—between 7% and 9% for different PAPA pairs— was close to that of nuclear SNAPf tag alone (7 %, Figure S2D and ref. ^53^), indicating that the level of observed P-TEFb:7SK chromatin binding is indistinguishable from background levels in our measurements. Moreover, a lower bound fraction was observed for SNAPf-CycT1 molecules reactivated by PAPA from HEXIM1-Halo (7.0 ± 0.6 %) and LARP7-Halo (7.8 ± 0.7 %) than for SNAPf-CycT1 molecules in the same cells reactivated by DR (13.6 ± 0.8% and 11.8 ± 0.7%, respectively). Thus, our findings imply that association of P-TEFb with 7SK inhibits chromatin binding and that most P-TEFb:7SK complexes are mobile in live cells.

### P-TEFb rapidly dissociates from 7SK upon Cdk9 inhibitor treatment

Prior work showed that P-TEFb is released from the 7SK complex when cells are treated with Cdk9 inhibitors. Using size-exclusion chromatography and immunoprecipitation of cell lysates, we likewise found that HEXIM1-SNAPf dissociates from LARP7-Halo after treatment of cells with the Cdk9 inhibitor NVP-2 (Figure S4A-B).^62^ A limitation of these methods is that they require lysis of cells followed by extended incubation periods, constraining their time resolution and potentially introducing artifacts by diluting proteins far below their endogenous concentrations. In principle, a perturbation that increases the K_d_ of a complex could leave it largely intact at endogenous protein concentrations, while causing it to dissociate when diluted in a lysate. We thus used PAPA to test whether NVP-2 causes P-TEFb to dissociate from 7SK in live cells and to measure the kinetics of this dissociation.

Using automated fSMT microscopy (see Methods), we made time course measurements of PAPA between the four sender-receiver pairs described above. NVP-2 triggered a rapid decline in PAPA signal in each case, with a half-decay time of around 3-5 minutes, and a final value close to estimated nonspecific background (Figures 3 and S4H). After 20 min of NVP-2 treatment, the diffusion profile of PAPA trajectories had become virtually identical to that of DR trajectories, further indicating that residual PAPA signal was nonspecific (compare Figure S4C-E to Figure 2D-F). Consistent with lysate-based experiments,^34,40^ HEXIM1 and CycT1 also dissociated—albeit more slowly—when cells were treated with the transcription inhibitor triptolide or with N,N’-hexamethalene bis-acetamide (HMBA), an inducer of HIV transcription (Figure S4F-H). Live-cell PAPA-fSMT measurements thus reveal that association of P-TEFb with 7SK can change drastically within minutes of an external stimulus.

**Figure 3:**
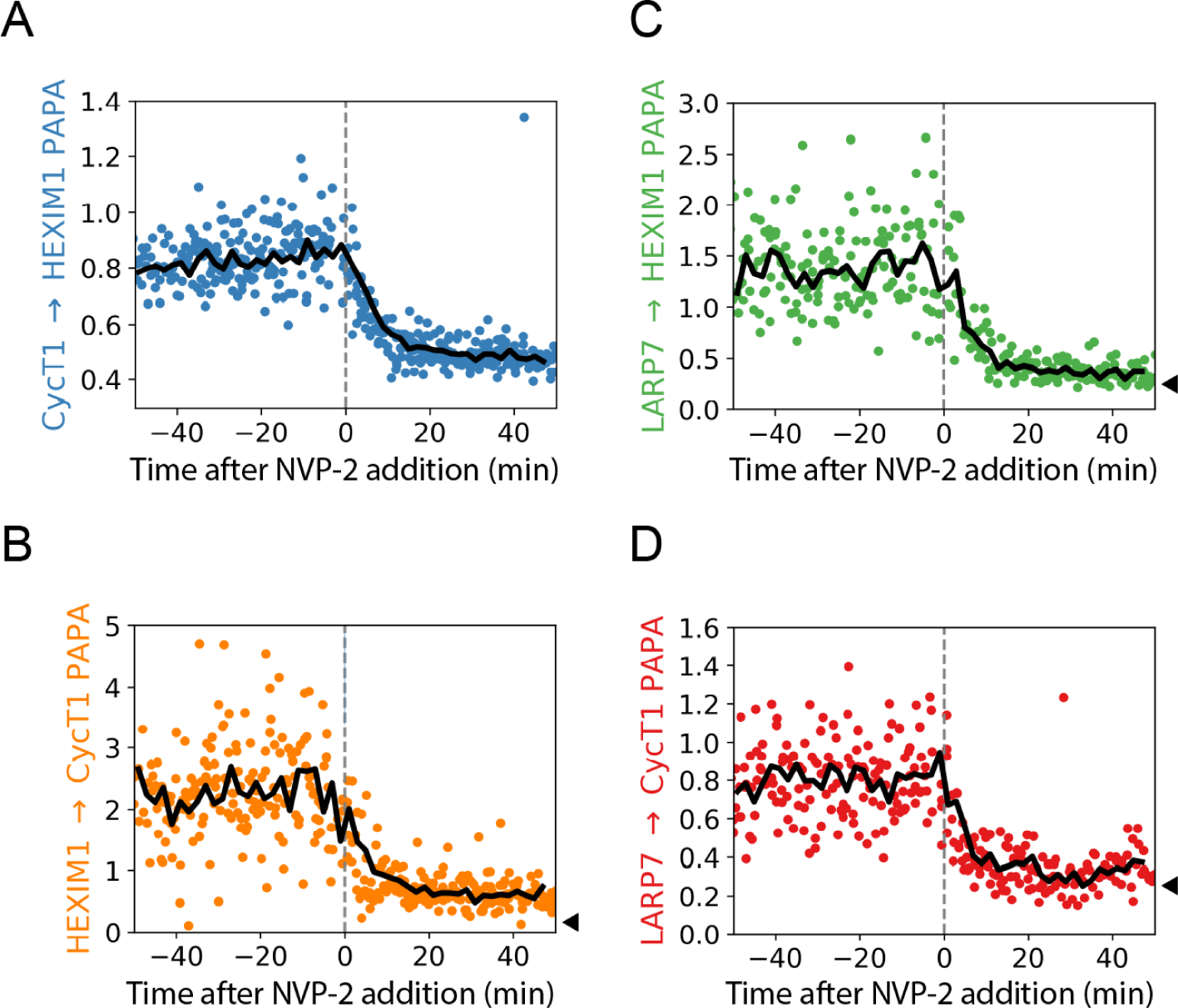
PAPA reveals rapid dissociation of the P-TEFb:7SK complex in response to the Cdk9 inhibitor NVP-2. As in Figure 2, each panel represents a different pair of sender- and receiver-labeled proteins. y-axes: normalized PAPA signal, calculated as the “GV ratio”—the increase in the number of localizations after green pulses (PAPA) divided by the increase after violet pulses (DR). x-axis: Time in minutes before or after addition of NVP-2. Each point represents a single movie, while solid lines represent averages over 2-min time bins. In each case, a rapid drop in PAPA signal was observed upon NVP-2 addition (dashed gray vertical line). Black carats on the right side of (B-D) represent background PAPA measurements between the corresponding Halo-tagged protein and H2B-SNAPf (see black lines in Figure S3A-B).

### Cdk9 inhibitor treatment rapidly alters CycT1 and HEXIM1 diffusion profiles

We predicted that if the P-TEFb:7SK complex diffuses at 2-3 µm^2^/s, then NVP-2 treatment should reduce the abundance of this subpopulation in fSMT measurements of CycT1 and HEXIM1. Indeed, the 2-3 µm^2^/s peak declined after NVP-2 treatment of HEXIM1-SNAPf cells, while the 6-7 µm^2^/s peak increased (Figure 4A). This shift was also apparent in cell-by-cell state array analysis (Figure S5A). While a similar shift was seen for SNAPf-CycT1 (Figures 4B and S5B), there remained a substantial fraction of molecules diffusing at 2-3 µm^2^/s, which might indicate the existence of other complexes of CycT1, not disrupted by NVP-2, that have a similar diffusion coefficient.

**Figure 4:**
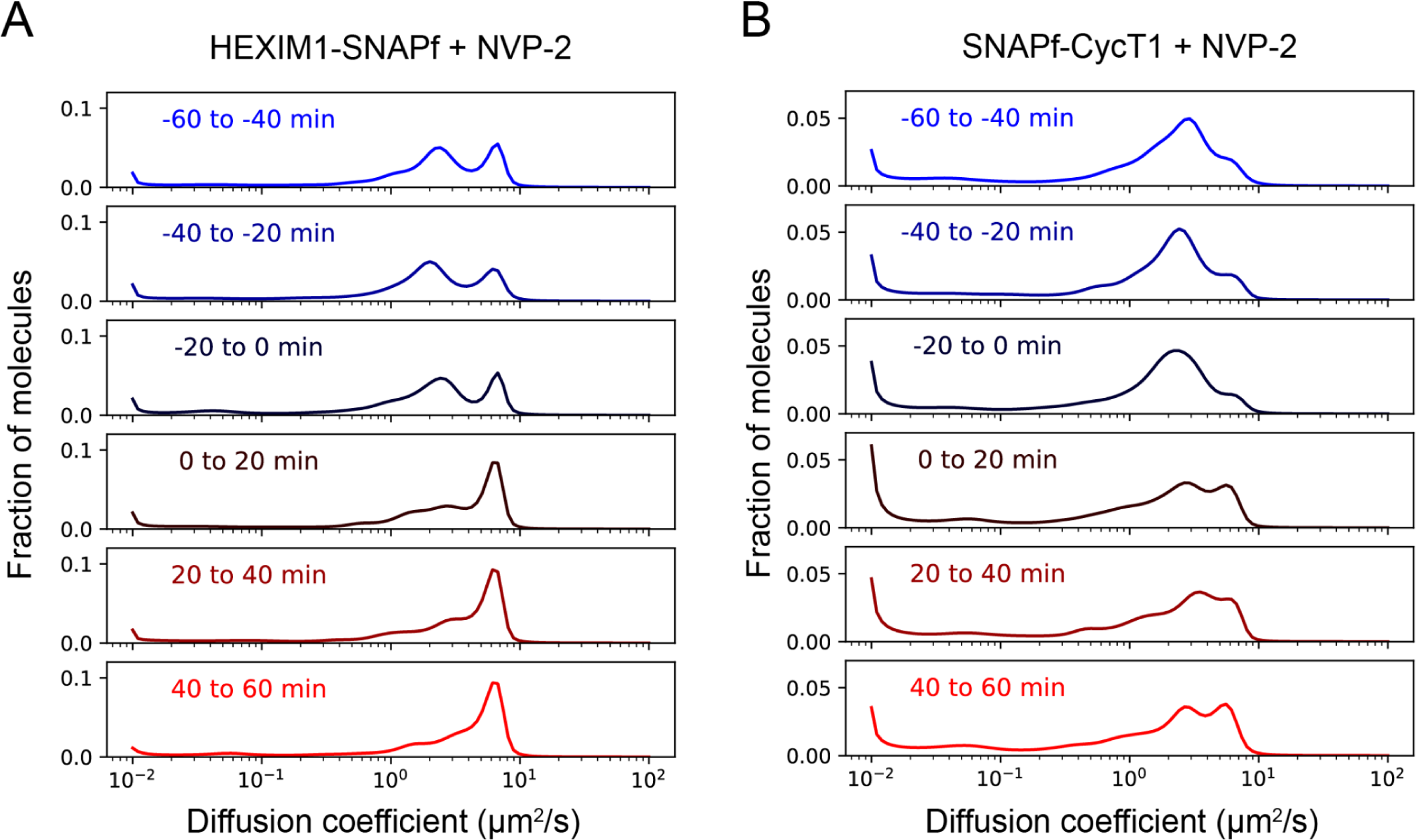
Single-component CycT1 and HEXIM1 diffusion profiles shift upon Cdk9 inhibition. Diffusion coefficient distributions of HEXIM1-SNAPf (A) and SNAPf-CycT1 (B) molecules as a function of time before or after addition of the Cdk9 inhibitor NVP-2. Tracking data from groups of cells were binned in 20-min time intervals prior to state array analysis. See Figure S5A-B for cell-by-cell state array analysis.

The immobile fraction of SNAPf-CycT1 (D < 0.1 µm^2^/s) increased slightly from 17% to a peak of 23% after NVP-2 addition (Figure S5D), consistent with relief of the inhibitory effect of 7SK on chromatin binding by P-TEFb. This increase was transient, however, as the bound fraction of CycT1 declined between 15 min and 2 hours after NVP-2 addition. The immobile fraction of HEXIM1 declined slightly from 11% to about 8% after NVP-2 treatment (Figure S5C).

Taken together, these results indicate rapid shifts in HEXIM1 and P-TEFb interactions in response to Cdk9 inhibitor treatment, consistent with PAPA-fSMT measurements of their dissociation from the 7SK complex.

### Monitoring hnRNP binding to the 7SK RNP complex

Cdk9 inhibitors have been proposed to release P-TEFb from 7SK indirectly by inhibiting transcription. According to this model, free hnRNPs—which ordinarily would have bound nascent transcripts—accumulate due to loss of mRNA synthesis and compete with P-TEFb for binding to 7SK. This model predicts that hnRNPs should interact with the 7SK complex in live cells and that this interaction should increase upon NVP-2 treatment.

To test whether different hnRNPs can interact with the 7SK complex in live cells, we performed PAPA-fSMT between endogenous LARP7-Halo and various transgenic SNAPf-tagged hnRNPs. A greater PAPA signal was observed for SNAPf-tagged hnRNP R (GV = 0.842 ± 0.019, 95% confidence interval), hnRNP Q (0.875 ± 0.031), hnRNP A1 (0.431 ± 0.010), and hnRNP H1 (0.318 ± 0.010) than for the H2B-SNAPf negative control (0.283 ± 0.006), consistent with previous affinity purification experiments in cell lysates (Figures 5B and S5E).^41,42,63^ In contrast, hnRNP C2, which was previously found not to interact with 7SK in immunoprecipitation experiments,^41^ showed only a background level of PAPA with LARP7 (0.278 ± 0.009; Figures 5B and S5E). Interaction with LARP7 was also detected for N-terminally SNAPf-tagged hnRNP Q, albeit with a lower PAPA signal (GV = 0.356 ± 0.010) than the C-terminally SNAPf-tagged protein (Figure S5F).

**Figure 5:**
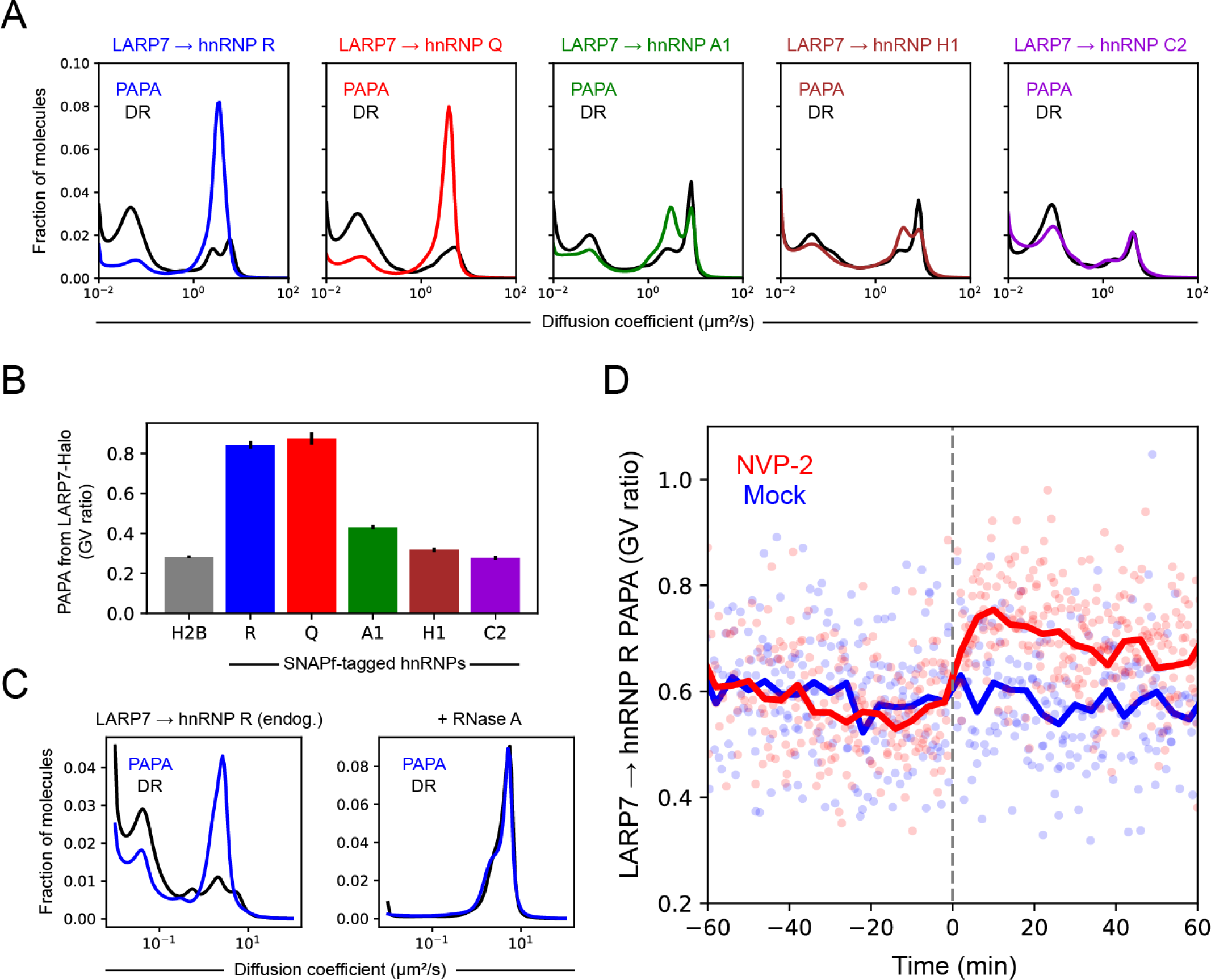
Interaction of hnRNPs with the 7SK complex. A-B) PAPA-fSMT analysis of stably transfected SNAPf-tagged hnRNPs in LARP7-Halo cells. A) Diffusion profiles of direct-reactivated molecules (DR, black curves) and molecules reactivated by PAPA from LARP7 (color curves). B) PAPA signal (GV ratio) between LARP7-Halo and each SNAPf-tagged hnRNP or H2B-SNAPf negative control (see also Figure S5E). C) Diffusion profiles of hnRNP R-SNAPf molecules reactivated by PAPA from LARP7-Halo (blue curves) and DR (black curves). Left panel shows cells bead-loaded with mNeonGreen-NLS protein alone, while right panel shows cells bead-loaded with mNeonGreen-NLS and RNase A. D) Effect of adding NVP-2 (red) or cell culture medium only (blue, “Mock”) on PAPA between endogenous LARP7 and hnRNP R. Points represent individual cells, while thick curves represent averages within 4-minute bins.

The diffusion coefficient distributions of each hnRNP exhibited a peak at the lowest diffusion coefficient in our state array, 0.01 µm^2^/s, as well as a slightly faster diffusing peak at 0.04-0.05 µm^2^/s (Figure 5A). Interestingly, this coincides with previous measurements of the diffusion coefficient of messenger ribonucleoprotein particles (mRNPs) in the nucleus, suggesting that it may represent hnRNPs bound to relatively long RNAs such as mRNAs.^64^ Finally, one or two distinct faster diffusing peaks were observed above 1 µm^2^/s (Figure 5A). For all hnRNPs with interaction signal above background, PAPA from LARP7-Halo enriched a peak at 3-4 µm^2^/s, indicating that the hnRNP:7SK complex resides within this peak (Figure 5A). Enrichment of the 3-4 µm^2^/s peak correlated with the GV ratios of the different hnRNPs, with strongest enrichment observed for hnRNPs R and Q, weaker enrichment for hnRNPs A1 and H1, and no enrichment for non-interacting hnRNP C2 (Figure 5A).

To further investigate which peaks in the hnRNP diffusion profile represent RNA-bound complexes, we repeated these measurements after bead-loading RNase A into a Halo-LARP7 + hnRNP R-SNAPf double homozygous knock-in cell line (Figure S2I). A purified fluorescent protein with a nuclear localization sequence (mNeonGreen-NLS) served as an internal control to identify bead-loaded cells. In contrast to cells bead-loaded with mNeonGreen-NLS alone, cells bead-loaded with mNeonGreen-NLS and RNase A retained only the fastest diffusing subpopulation of hnRNP R (Figures 5C). RNase A reduced PAPA signal to the level of nonspecific background (Figure S5G), indicating that association between LARP7 and hnRNP R is mediated by RNA. Moreover, RNase A caused the diffusion profile of PAPA-reactivated hnRNP R-SNAPf molecules to become indistinguishable from that of direct-reactivated molecules (Figure 5C, right panel), consistent with the residual PAPA signal being nonspecific. These results suggest that the fastest peak in the diffusion profile of hnRNP R represents protein not bound to RNA, while the slower peaks represent complexes of hnRNP R with 7SK and other RNAs.

Finally, to test whether release of P-TEFb from the 7SK complex coincides with binding of hnRNPs, we monitored PAPA between endogenous LARP7 and hnRNP R upon NVP-2 treatment. LARP7 → hnRNP R PAPA increased after NVP-2 treatment (Figure 5D; see also Figure S6 and Methods) with kinetics similar to, if not slightly faster than P-TEFb dissociation from 7SK (Figure S4H). These results support the idea that Cdk9 inhibition causes 7SK to exchange partners, releasing P-TEFb while binding hnRNPs.

### A viral activator dissociates P-TEFb from HEXIM1

The HIV transcriptional activator Tat binds P-TEFb and tethers it to nascent viral transcripts to release Pol II from pausing. Transient transfection and *in vitro* biochemical experiments have shown that Tat can displace P-TEFb and HEXIM1 from 7SK.^37,39,65^ To test whether this displacement is detectable by PAPA in live cells, we transfected HEXIM1-Halo + SNAPf-CycT1 cells with either mNeonGreen-labeled Tat or mNeonGreen with a nuclear localization sequence (NLS) as a negative control. We measured the HEXIM1 → CycT1 PAPA signal, along with the fluorescence intensity in the mNeonGreen channel, across many cells. No correlation was observed between HEXIM1 → CycT1 PAPA signal and mNeonGreen-NLS intensity (Spearman ⍴ = 0.06, p = 0.27, Figure 6A). However, a strong negative correlation was observed between HEXIM1 → CycT1 PAPA and Tat-mNeonGreen intensity (⍴ = −0.65, p = 3 × 10^-88^, Figure 6A), consistent with Tat-induced dissociation of P-TEFb from HEXIM1. Thus, PAPA provides a way to measure release from inhibition of endogenous P-TEFb by a transcriptional activator in live cells.

**Fig. 6:**
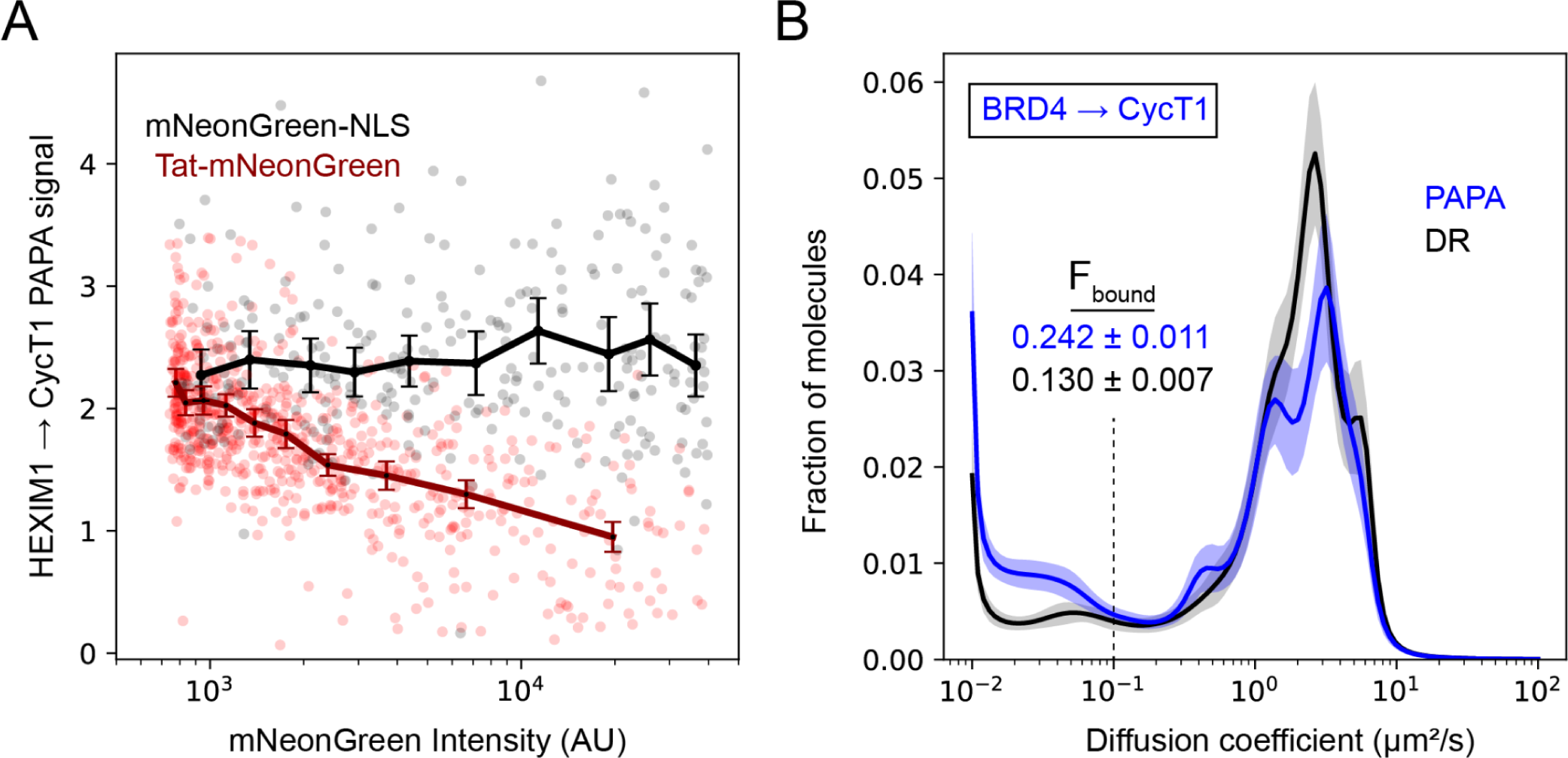
Influence of transcriptional activators on P-TEFb. A) Concentration-dependent dissociation of the PTEFb:7SK complex by the HIV transcriptional activator Tat. PAPA signal (GV ratio) in HEXIM1-Halo + SNAPf-CycT1 cells transiently transfected with expression plasmids encoding Tat-mNeonGreen or mNeonGreen-NLS negative control. Transparent points represent individual cells, while solid lines with error bars represent mean ± 2*S.E. of sets of cells binned by decile of mNeonGreen intensity. B) State array analysis of CycT1-SNAPf reactivated by direct reactivation (DR, black) or PAPA from Halo-BRD4 (blue). Solid curves and transparent regions represent the mean and approximate 95% confidence intervals (± 2 * S.D.) from bootstrapping analysis. Dashed gray line represents the D = 0.1 µm^2^/s cutoff for calculating the fraction bound to chromatin.

### Binding of P-TEFb to BRD4 correlates with binding to chromatin

P-TEFb also binds the endogenous transcriptional coactivator BRD4.^7,66–69^ To assay this interaction in live cells, we performed PAPA-fSMT measurements in a Halo-BRD4 + SNAPf-CycT1 double knock-in cell line. BRD4 → CycT1 PAPA signal was significantly greater than a BRD4 → SNAPf negative control (GV ratio 0.546 ± 0.015 vs. 0.221 ± 0.013; Figure S5H), indicating specific association between P-TEFb and BRD4. State array analysis revealed that PAPA trajectories had a substantially higher bound fraction (24.2 ± 1.1%) than DR trajectories (13.0 ± 0.7%, Figure 6B). This implies that association of P-TEFb with BRD4 positively correlates with chromatin binding, consistent with a model in which BRD4 recruits P-TEFb to chromatin to activate target genes.

## Discussion

Transcription, like other cellular processes, is regulated by extensive networks of interactions between macromolecules. Accurately defining these networks and illuminating their regulatory mechanisms will require studying them in live cells, where the various components, and their competing partners, are organized in the appropriate subcellular compartments at physiological concentrations. Because regulation hinges on the rates of these interactions, it will moreover be essential to measure how fast molecular complexes form and dissociate.

To meet this challenge, our field will need to develop new experimental methods. Conventional biochemical approaches requiring cell lysis massively dilute molecular complexes and obliterate their spatial organization. Fixation-based methods freeze interactions in place, obscuring their dynamics, and biasing our measurements toward interactions that are relatively stable and easier to crosslink. Fluorescence imaging methods like fSMT overcome some of these limitations, yet our ability to monitor molecular interactions has remained limited.

Proximity-assisted photoactivation (PAPA), in combination with fSMT, provides a promising new strategy to study molecular interactions in live cells, while preserving the physiological concentrations, spatial organization, and interaction kinetics of endogenous macromolecules. Here we have described two key advances in the development and application of this approach.

First, we have combined PAPA with fSMT to visualize specific endogenous complexes. Fast SMT of fluorescently labeled CycT1 and HEXIM1, followed by state array analysis, revealed at least two distinct mobile subpopulations of each protein with different diffusion coefficients (Figures 2A-B and S1B-C). PAPA between four different sender- and receiver-labeled protein pairs showed that the P-TEFb:7SK complex resides within the slower-diffusing subpopulation (2-3 µm^2^/s) (Figure 2C-F), consistent with the much greater molecular mass of this complex than either P-TEFb or HEXIM1 alone. These measurements reveal that most P-TEFb:7SK complexes are not bound to chromatin, an important feature that previous lysis- and fixation-based experiments had left unresolved. Fast SMT measurements similarly revealed multiple subpopulations of hnRNPs with distinct mobilities, and PAPA indicated that the hnRNP:7SK complex resides within one of these subpopulations (D = 3-4 µm^2^/s). Our results are consistent with previous in vitro evidence showing that multiple hnRNPs bind 7SK. We anticipate that PAPA-fSMT will be useful more broadly to probe the diverse interaction networks between RNA binding proteins and RNAs involved in transcription, splicing, and translation.

Second, we were able to detect the exchange of subunits within endogenous regulatory complexes in live cells in real time (Figure 7). While previous cell lysis and biochemical fractionation experiments estimated that P-TEFb dissociates from 7SK within 15-30 min of a perturbation,^23,70^ PAPA revealed that this dissociation occurs within several minutes (Figure 3). Synchronous loss of PAPA for all four pairs of proteins tested implies that HEXIM1 and P-TEFb are released from the 7SK complex simultaneously within the time resolution of our measurements (Figure S4H). At the same time, hnRNP R binding to 7SK increased with similar kinetics after Cdk9 inhibitor treatment (Figures 5D and S4H), consistent with a model in which hnRNPs immediately replace P-TEFb in the 7SK RNP complex.

**Figure 7:**
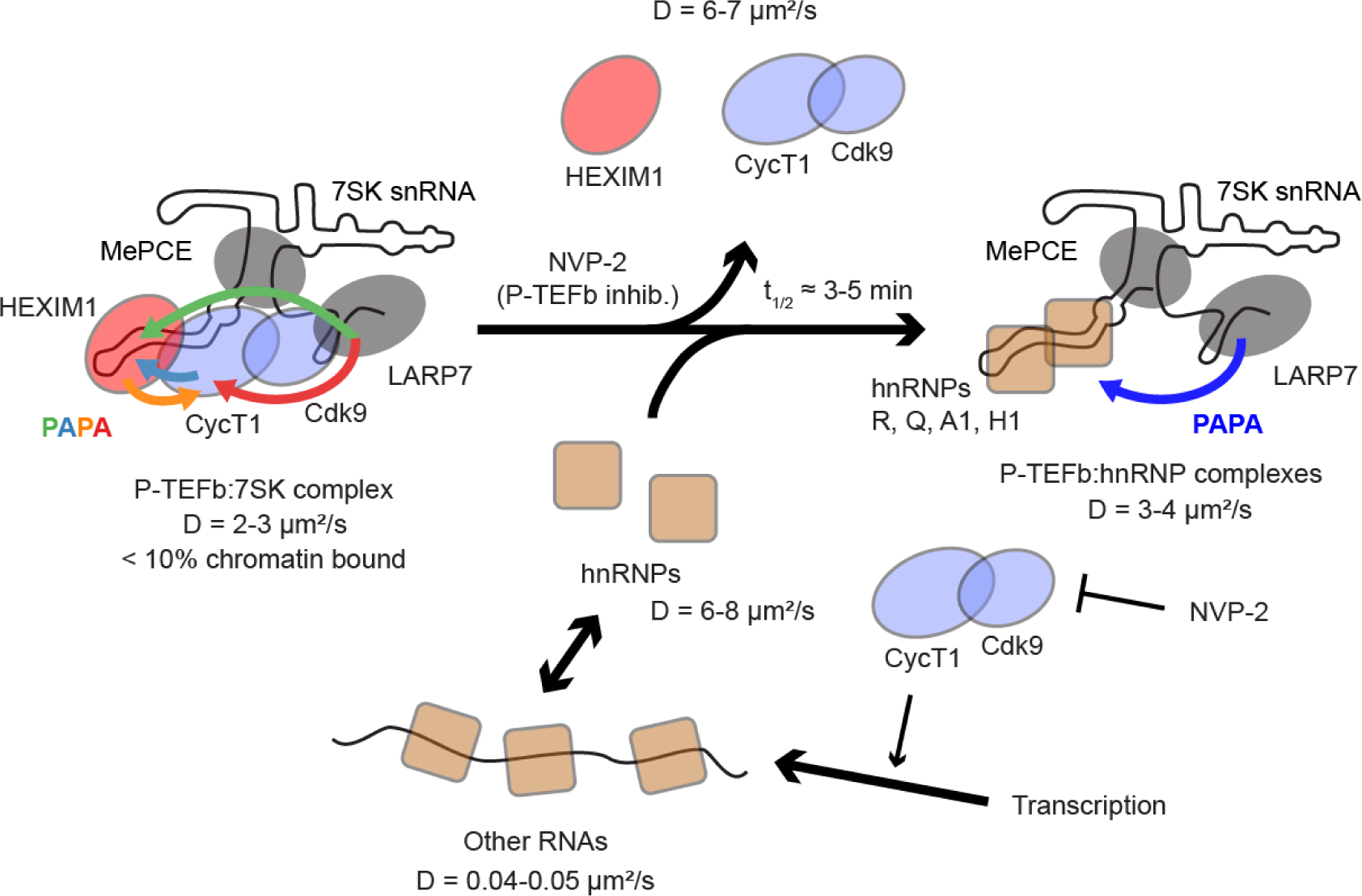
Visualizing endogenous P-TEFb regulatory complexes using SMT and PAPA. PAPA-SMT between four pairs of endogenous proteins revealed that the P-TEFb:7SK complex is predominantly unbound to chromatin and diffuses with a characteristic diffusion coefficient of 2-3 µm^2^/s. P-TEFb dissociates from the 7SK complex within a several minutes of treatment with NVP-2 and is replaced with hnRNPs such as hnRNP R.

We also observed the exchange of binding partners of P-TEFb. Expression of one such partner, the HIV transcriptional activator Tat, disrupted the association of P-TEFb with the 7SK complex in a dose-dependent manner (Figure 6A). PAPA also detected interaction of P-TEFb with the endogenous coactivator BRD4 (Figures 6B and S5H). While 7SK binding was associated with reduced chromatin occupancy of P-TEFb (Figure 2D,F), BRD4 binding was associated with increased chromatin occupancy, consistent with the proposed role of BRD4 in recruiting P-TEFb to target genes (Figure 6B).

Our work raises multiple questions about P-TEFb regulation. First, it is unclear why PAPA-fSMT detected minimal interaction of P-TEFb:7SK with chromatin, in contrast to the widespread interactions suggested by crosslinking-based methods. While a trivial explanation is that crosslinking is prone to artifacts, another possibility is that P-TEFb:7SK may bind chromatin infrequently or transiently before dissociating to release P-TEFb.^45^

Second, our results support the model that hnRNPs replace P-TEFb in the 7SK complex, yet it remains unclear whether this competition is regulated solely by the concentration of free hnRNPs and mass-action kinetics. Given the rapid and complete dissociation of P-TEFb from 7SK that we observe in response to a Cdk9 inhibitor, a mass action mechanism would seem to require a correspondingly rapid fold-change in the concentration of free hnRNPs to displace P-TEFb from 7SK. This in turn would imply that at steady-state, available binding sites on RNA are largely occupied by hnRNP molecules, such that transcription inhibition would lead to rapid saturation of available sites and accumulation of excess free hnRNPs. In contrast, we observe a substantial population of fast-diffusing hnRNPs that are presumably unbound to RNA (Figure 5A,C)—with the caveat that this could be an artifact of protein overexpression (Figure 5A) or degradation (Figures 5C and S2I). Taken together, our observations raise the question whether P-TEFb release from 7SK is regulated by more direct and rapid mechanisms than hnRNP accumulation.

Third, it is unclear why binding of P-TEFb to chromatin transiently increases and then decreases upon treatment with a Cdk9 inhibitor (Figure S5D). One hypothesis is that this reflects interaction of P-TEFb with elongating polymerases, which finish transcribing but are not replenished while pause release is blocked by Cdk9 inhibition.^71^ An important goal for future PAPA-fSMT experiments will be to delineate which molecular complexes comprise the “chromatin-bound” fraction of P-TEFb.

Fourth, it is unclear how many other distinct P-TEFb complexes are represented in the mobile sub-population of CycT1 and how interaction of P-TEFb with these complexes is regulated. Given the diversity of other complexes predicted to bind P-TEFb,^45,72–75^ and the complicated shift that we observed in the CycT1 diffusion profile upon NVP-2 treatment (Figure 4), we suspect that we have only scratched the surface of the P-TEFb interactome. PAPA-fSMT with other partners could help to tease apart this extended network of competing and multifunctional interactions.

More broadly, we expect that the PAPA-fSMT approaches that we have developed to study P-TEFb regulation will be applicable to many other cellular processes. Advances in molecular biology have historically been driven by the development of new technologies to detect, separate, and characterize macromolecular complexes. PAPA and fSMT extend this biochemical toolkit into live cells. While the complexes that we focused on here are relatively stable, cells contain vast networks of weak, ephemeral molecular interactions that have proven difficult, if not impossible to reconstitute in vitro, preserve in lysates, or pin down by crosslinking. PAPA-fSMT will make it possible to examine such interactions in their native context.

### Limitations of the study

Multiple complexes with overlapping diffusion coefficients may blur into a single peak in state array analysis. We suspect that this may be true for CycT1, which is known to bind many partners, and whose diffusion profile shifted in a complicated way upon treatment with NVP-2 (Figure 4B). Diffusion coefficients of different complexes might be better resolved in the future by tracking more molecules or by tracking molecules with higher time resolution.^76,77^

Fusion to Halo and SNAPf tags sometimes disrupts protein function and interactions. While both Halo- and SNAPf-tagged CycT1 activated a reporter gene as expected (Figure S1A), we observed a difference in the shape of their diffusion profiles (Figure S1B), suggesting that one tag or both may shift the distribution of P-TEFb between different complexes.

In addition to reactivating specific complexes, PAPA can nonspecifically reactivate receiver-labeled molecules that come close to a sender-labeled molecule by random chance. While experiments with negative control proteins let us estimate this background reactivation and distinguish it from true association (Figures S3A-B and S5E-H), an important future goal will be to develop analysis methods to infer more accurately the diffusion profile of complexes in the presence of nonspecific background.

Our current measurements of PAPA efficiency (GV ratios) are in relative units, which does not tell us what fraction of a molecule interacts with a partner. We cannot say, for instance, whether the higher PAPA signal from LARP7 to hnRNPs R than to hnRNP A1 reflects greater specificity of hnRNP R for 7SK or closer proximity of Halo and SNAPf tags within the LARP7:7SK:hnRNP R complex. Future method development will be required to deconvolve these contributions to the observed PAPA efficiency.

## Acknowledgments

Thanks to John Lis, Philip Versluis, Koh Fujinaga, Matija Peterlin, and members of the Tjian-Darzacq group for helpful discussions. HIV-luciferase reporter and Myc-Tat expression plasmids were generously provided by Koh Fujinaga. Sathvik Anantakrishnan generously shared code for cell segmentation in automated imaging, and Nike Walther provided valuable feedback for the refinement of automated fSMT acquisition and analysis methods. This work was supported by the Chan-Zuckerberg Initiative Dynamic Imaging grant (to X.D.). R.T. is an Investigator of the Howard Hughes Medical Institute.

## Methods

### Cell line maintenance and genetic manipulation

U2OS cells were maintained in Dulbecco’s modified Eagle’s medium (DMEM) with 4.5 g/l glucose (Thermo Fisher #10566016), 10% fetal bovine serum (FBS), and 100 U/ml penicillin-streptomycin (Thermo Fisher #15140122). Cells were routinely tested for mycoplasma contamination by PCR.

For genome editing, Fugene 6 was used to co-transfect cells with a homology donor plasmid together with a second plasmid encoding SpCas9, the fluorescent protein Venus, and an sgRNA gene driven by a U6 promoter. In some cases, Venus-expressing cells were bulk sorted after one week, cultured an additional week, and then single-cell sorted into 96-well plates, using staining with fluorescent Halo or SNAP ligands to enrich for edited cells. In other cases, single-cell sorting of Halo/SNAP ligand-labeled cells was performed after one week without a preceding Venus-sorting step. We have come to prefer the latter method.

Single-cell clones were re-plated in a duplicate 96-well plate, and DNA was extracted from each clone by 15 min incubation in a homemade lysis buffer (10 mM Tris pH 7.5, 1 mM CaCl_2_, 3 mM MgCl_2_, 1 mM EDTA, 1% Triton X-100, 250 µg/ml proteinase K). After heat-inactivation of proteinase K for 1 hour at 85°C, correct insertion of tags was confirmed by PCR with primers outside the homology arms, followed by Sanger sequencing of PCR products.

PiggyBac transgenic lines were generated by co-transfecting cells with transposon and transposase plasmids and then adding puromycin after two days to a final concentration of 1 µg/ml to select for integrants. Puromycin selection was continued for at least one week before performing experiments.

### Preparation of cells for fSMT and PAPA-fSMT experiments

Cells were trypsinized, counted using a Countess automated cell counter (Thermo Fisher), centrifuged, and resuspended in phenol red-free DMEM (Thermo Fisher #21063029) supplemented with 10% fetal bovine serum (FBS), 100 U/ml penicillin-streptomycin (Thermo Fisher #15140122), and GlutaMax glutamine supplement (Thermo Fisher #35050061). Cells were plated at a density of 400,000 cells per dish in 2 ml of phenol red-free DMEM in 35-mm glass-bottom dishes (MatTek P35G-1.5-20-C).

### Automated microscopy

We engineered an automated system to perform fSMT measurements with higher throughput (see also refs. ^53,54^). Using code written in the Nikon Elements macro language and Python (https://github.com/tgwgraham/autosmt_v4tg2), we programmed a Nikon Ti-E microscope to raster-scan a rectangular grid of different fields of view in a sample, which were spaced far enough apart to avoid photobleaching of adjacent fields of view. As a further precaution to avoid exposure of subsequent fields of view to laser light, the scan direction was chosen to be opposite to that of the oblique laser beam used for HILO illumination. At each grid position, a snapshot image was acquired, and nuclei were identified using the StarDist segmentation package. The stage was repositioned to approximately center a target nucleus in the field of view, a second snapshot image was acquired and used to re-locate the target nucleus, and an fSMT movie was acquired within a rectangular region of interest (ROI) circumscribed on the nuclear mask.

The power densities of each laser at the sample, with the acousto-optic tunable filter (AOTF) set to 100%, were approximately 67 W/cm^2^ for 405 nm, 190 W/cm^2^ for 561 nm at the power level used for green light-induced reactivation in PAPA experiments, 2.2 kW/cm^2^ for the full-power setting on the 561 nm laser used for SMT imaging of JFX549, and 2.3 kW/cm^2^ for 639 nm. Power density was measured by using an iris to shrink the illuminated region to fit inside the camera field of view, measuring the laser power passing through the objective with a laser power meter (Thorlabs PM100D and S170C), and dividing this value by the illuminated area.

### fSMT experiments

Because different endogenous proteins have different concentrations (Figure S2), staining and imaging conditions were optimized empirically for each target protein to obtain a sufficient sample size of trajectories while limiting molecular localizations to a trackable density.

To track SNAPf-tagged proteins by direct reactivation, cells were stained overnight with 5 nM JFX650 SNAP ligand (Janelia Materials) in phenol red-free DMEM, rinsed twice with phosphate-buffered saline (PBS), destained twice for 30 min in phenol red-free DMEM, and exchanged into fresh medium once more before imaging. Each cell was pre-bleached for 10 s with full power using a 639 nm laser. Three 2000-frame intervals were collected at 7.48 ms/frame with 2 ms stroboscopic full-power 639 nm illumination and 2%, 4%, and 6% (or in some cases, 10%) acousto-optic tunable filter (AOTF) power on the 405 nm laser in the inter-frame interval. This stepwise increase in 405 nm laser power served to counteract the gradual depletion of dark-state molecules due to reactivation and photobleaching.

To track Halo-CycT1 molecules by direct reactivation, cells were labeled overnight with 50 nM JFX549 Halo ligand in phenol red-free DMEM, rinsed twice with PBS, destained twice for 30 min, and exchanged into fresh medium once more before imaging. Each cell was pre-bleached for 5 s with full power using a 561 nm laser. Three 1000-frame intervals were collected at 7.48 ms/frame with 2 ms stroboscopic full-power 561 nm illumination and incrementally increasing 2, 4, or 6% AOTF power on the 405 nm laser in the inter-frame interval. A similar protocol was used for dSTORM of HEXIM1-Halo, but with 15 min staining with 10 nM JFX650 HTL and 15 min destaining, 639 nm rather than 561 nm illumination, a 10 s prebleaching period, and 2000 frames in each interval.

For SMT of LARP7-Halo (Figure S3C), cells were labeled for 20-30 min with a mixture of 50 nM JFX549 Halo ligand and 100 pM JFX650 Halo ligand. Cells were located in the densely labeled JFX549 channel and imaged in the sparsely labeled JFX650 channel, for 500 frames at 7.48 ms/frame with 2 ms stroboscopic full-power 639 nm illumination.

### PAPA experiments

For PAPA experiments, cells were incubated with 50 nM JFX549 Halo ligand and 5 nM JFX650 SNAP ligand overnight. Prior to imaging, cells were rinsed twice with PBS, destained twice for 30 min in phenol red-free DMEM, and exchanged once more into fresh medium.

Cells were located using snapshot images taken in the JFX650 channel with red (639 nm) light at 2-5% power. After centering the cell in the field of view and taking snapshot images in both the JFX549 and JFX650 channels, JFX650 was pre-bleached for either 5 or 10 seconds with full-power red light, and the selected cell was imaged in a cropped field of view with 5 repetitions of the following illumination sequence:

1. Bleaching: 1.4 seconds of red light, not recorded
2. Pre-DR imaging: 30 frames of red light, 2 ms stroboscopic pulse per frame
3. DR: violet (405 nm) light pulse, not recorded
4. Post-DR imaging: 30 frames of red light, 2 ms stroboscopic pulse per frame
5. Bleaching: 1.4 seconds of red light, not recorded
6. Pre-PAPA imaging: 30 frames of red light, 2 ms stroboscopic pulse per frame
7. PAPA: green (561 nm) light pulse, not recorded
8. Post-PAPA imaging: 30 frames of red light, 2 ms stroboscopic pulse per frame

This illumination sequence is optimized from our initial protocol^55^ and is similar to that described elsewhere.^53^ It collects fewer total frames to reduce file sizes and includes 1.4-second (200-frame) periods of continuous red illumination (steps 1 and 5) to bleach reactivated molecules and thereby minimize cross-contamination of DR and PAPA trajectories.

Because of differences in the protein concentrations and PAPA efficiencies of different protein pairs, the duration of green and violet pulses was adjusted empirically between different cell lines, as indicated in the table below, to achieve a density of localizations high enough to acquire reasonable sample sizes yet sparse enough to track. Importantly, for a given Halo-tagged, sender-labeled protein, the same conditions were used to compare SNAPf-tagged, receiver-labeled interactors with non-interacting controls.

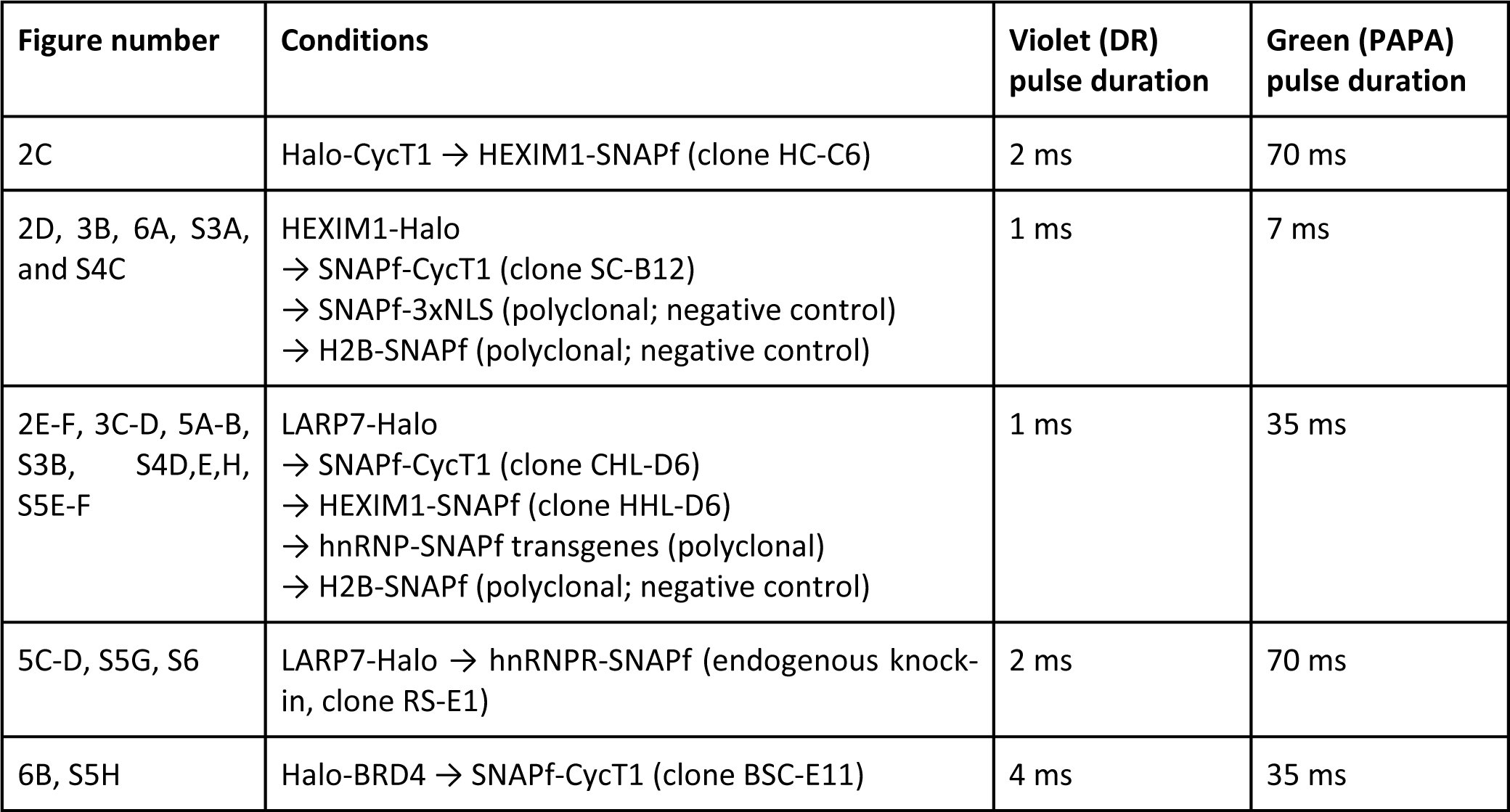

The CycT1 → HEXIM1 timecourse experiments in Figures 3A and S4F-H employed an earlier version of this protocol, listed below, in which cells were pre-bleached with 8.65 s of full-power 639 nm light, the full 512 x 512 pixel field of view was imaged at a frame rate of 18 ms/frame, and localizations were counted over the entire field of view.

1. Bleaching: 1.35 seconds of red (639 nm) light, not recorded
2. Pre-PAPA imaging: 40 frames of red light, 2 ms stroboscopic pulse per frame
3. PAPA: green (561 nm) light pulse, 20 frames (360 ms), not recorded
4. Post-PAPA imaging: 40 frames of red light, 2 ms stroboscopic pulse per frame
5. Bleaching: 1.35 seconds of red (639 nm) light, not recorded
6. Pre-DR imaging: 40 frames of red light, 2 ms stroboscopic pulse per frame
7. DR: violet (405 nm) light pulse, 4 ms, not recorded
8. Post-DR imaging: 40 frames of red light, 2 ms stroboscopic pulse per frame

### fSMT and PAPA data analysis

Molecules were localized and tracked using quot (https://github.com/alecheckert/quot), and trajectories were analyzed using state array analysis (https://github.com/alecheckert/saspt). A range of diffusion coefficients between 0.01 µm^2^/s and 100 µm^2^/s was used for the state array, which is predicted to encompass the dynamic range of diffusion coefficients resolvable given our acquisition and tracking settings.^77^ As a quality-control step, a home-built “cellpicker” utility written in MATLAB (https://github.com/tgwgraham/basic_PAPASMT_analysis) was used to review images of cells from automated experiments and select cells for analysis. Cells were excluded if they appeared dead, misshapen, or abnormally bright or dim. For PAPA experiments, trajectories were pooled from the 30 stroboscopic 639 nm imaging frames after the 561 nm laser pulse (“PAPA trajectories”) or after the 405 nm laser pulse (“DR trajectories”), and state array analysis was separately performed for each set of trajectories. Analysis code can be found on GitHub (https://github.com/tgwgraham/basic_dSTORM_analysis, https://github.com/tgwgraham/basic_PAPASMT_analysis). GPT-3 (OpenAI) was used as a coding aid.

Confidence intervals for diffusion coefficient histograms were estimated by applying state array analysis to 100 bootstrap replicates randomly resampled by cell with replacement. To reduce computation time, the analysis was limited to 30,000 randomly selected trajectories per replicate. Shaded regions in Figures 2C-F and 6B represent the mean of all replicates plus or minus two standard deviations of the bootstrap distribution.

The single-cell green-to-violet (GV) ratios plotted in Figures 3, 5D, 6A, S4F-H, and S6 were calculated by dividing the difference in the total number of single-molecule localizations before and after green pulses (PAPA) by the difference before and after violet pulses (DR). Cells were excluded that had fewer than 500 total violet-reactivated localizations over the entire movie. To avoid artifacts due to camera saturation in Figure 6A, cells were excluded if the mean intensity in the mNeonGreen channel exceeded 40000 counts. For unknown reasons, an initial transient decrease in PAPA signal was observed in LARP7 → hnRNP R time course experiments (Figure S6), and the first 30 time points in each time course are excluded from the plot in Figure 5D.

Ensemble GV ratios were determined by fitting the difference in total number of localizations due to green and violet pulses to a line passing through the origin (Figures S3A-B and S5E-H). 95% confidence intervals were calculated as the fitted slope plus or minus the square root of the variance in the fit times the t-critical value for alpha = 0.05.

### Drug-addition experiments

NVP-2 (Cayman #34725, CAS #1263373-43-8) was used for all experiments at a final concentration of 400 nM. Triptolide (Selleckchem #S3604) was used at a final concentration of 3.3 µM, and N,N’-Hexamethylene bis(acetamide) (HMBA; Sigma 224235) was used at a final concentration of 10 mM [sic], as in previous work.^34,51,78,80^ In some experiments, the drug was dissolved at 3 times the final concentration in 1 ml of medium and added manually to 2 ml of medium in a MatTek dish. After mixing well by pipetting (8-10 times), 1 ml of medium was withdrawn to maintain the same volume of liquid in the dish. A 2-piece custom 3D printed lid, along with a magnetic clamp, was used to hold the dish in place while adding the drug (https://github.com/tgwgraham/dish_lids_35mm/). In other experiments, we used a home-built automated injection system: A syringe pump (Aitoserlea, Amazon) was controlled by an Arduino microcontroller, which in turn was triggered by a Python script via a serial connection from the microscope computer. The Python script was executed from the NIS Elements macro after imaging a specified number of fields of view. Compounds diluted at 2 times the final concentration in 2 ml of medium were injected from a 5 ml syringe via PE60 polyethylene tubing (Intramedic 427416), which was connected to a 3D printed MatTek dish lid fitted with a metal tube made by cutting a 90°-bent, blunt-ended 21-gauge needle. The pump was programmed to mix the solution by withdrawing and dispensing 2 ml of liquid five times. Drawings and source code for this setup are available at https://github.com/tgwgraham/autoinjector.

### Dual luciferase assay

Cells were seeded in 12-well plates at a density of 200,000 per well and transfected using 1.5 µl/well FuGENE 6 (Promega) with 5 ng of CMV-Renilla plasmid (pTG888), 250 ng of HIV-luc reporter (pTG1044), and 250 ng of the indicated Tat and mNeonGreen expression plasmids (pTG1033, pTG1035, pTG1036, or pTG1043; see Table S3: List of Plasmids and https://github.com/tgwgraham/ptefb_papa_plasmids). The next day, cells were analyzed using the Dual-Luciferase® Reporter Assay System (Promega #E1960). Culture medium was aspirated, and cells were rinsed once with PBS and lysed in 250 µl of passive lysis buffer for 15 min at room temperature on a rocker. Lysates were analyzed for firefly and *Renilla* luciferase activities on a GloMax® 20/20 luminometer (Promega), following the manufacturer’s instructions.

### Fluorescent gels and western blots

Cells were stained overnight with 50 nM JF549/JFX650 SNAP ligand and 50 nM JFX650/JFX549 Halo ligand, and lysates were prepared as previously described.^55,79^ A quantity of each lysate equivalent to 100,000 cells was loaded on a 10% Bis-Tris SDS-PAGE gel cast using custom 3D printed gel combs (https://github.com/tgwgraham/gel_combs). JFX549/JF549 and JFX650 were imaged on a Pharos FX Plus Molecular Imager (BioRad) using the “Low Sample Intensity” setting in the Cy3 and Cy5 channels. Molecular weights were determined using Precision Plus Protein All Blue Prestained Protein Standards (BioRad #1610373), which fluoresce in the Cy5 channel.

For western blotting, proteins were transferred to nitrocellulose membranes (Amersham #10600041), which were blocked for 30 min in 5% bovine serum albumin (BSA) in Tris-buffered saline with Tween-20 (TBST; 10 mM Tris-HCl, pH 7.4, 500 mM NaCl, 0.1% Tween-20). For Figure S2C,E, membranes were probed for approximately 3 h with 1:1000 dilutions of goat anti-CycT1 (Santa Cruz #sc-8127) or rabbit anti-HEXIM1 (Bethyl A303-112A), washed three times for 5 min with TBST, re-probed for 1 hour with 1:5000 HRP-conjugated secondary antibodies (donkey anti-goat, Santa Cruz #sc-2020; goat anti-rabbit, Invitrogen #31462), washed three times for 5 min with TBST, and imaged on a ChemiDoc MP (Bio-Rad) using Western Lightning chemiluminescent HRP substrate (Perkin Elmer #NEL105001EA). For Figure S2D,F, previously blotted membranes from Figure S2C,E were re-probed with 1:5000 mouse monoclonal anti-TATA binding protein antibody (Abcam #ab51841). The membrane in Figure S2H was probed for 1 h with 1:1000 rabbit anti-LARP7 antibody (Fortis/Bethyl #A303-723A) in 5% BSA/TBST, washed three times for 15 min with TBST, probed overnight with 1:5000 HRP-conjugated goat anti-rabbit antibody, washed three times for 15 min with TBST, and imaged as above.

### Immunoprecipitation and size-exclusion chromatography of cell lysates

Whole-cell lysates were prepared using an modification of a previous protocol.^41^ Confluent cells in 10-cm dishes were stained overnight with 50 nM JFX549 Halo ligand and 50 nM JFX650 SNAP ligand. Dishes were treated for 30 min with medium supplemented with either 400 nM NVP-2 or solvent only (ethanol) as a negative control. Cells were washed twice with ∼10 ml of PBS and once with 2.5 ml of wash buffer (10 mM Na-HEPES pH 7.9, 1.5 mM MgCl_2_, 10 mM KCl, 200 mM NaCl, 0.2 mM EDTA). After thoroughly aspirating the wash buffer, cells were scraped off the dish and resuspended by pipetting in 1 ml of lysis buffer (wash buffer supplemented with 1 mM DTT, 1 mM PMSF, 0.5% NP-40 alternative (Calbiochem), and one EDTA-free cOmplete protease inhibitor cocktail tablet [Roche] per 10 ml). The cell suspensions were left at room temperature for 10 min with periodic inversion and were then centrifuged for 5 min at maximum speed in a 4°C microcentrifuge. The soluble supernatant was collected, and the insoluble pellet was resuspended for SDS-PAGE analysis in 900 µl of lysis buffer with 300 µl of 4x SDS loading buffer without dye (see above).

For immunoprecipitation (Figure S4A), 1 ml of soluble lysate was incubated for 30 min at room temperature on a rotator with 40 µl bed volume of anti-Flag M2 agarose beads (Sigma) that had been pre-washed twice with the same buffer. Tubes were centrifuged for ∼30 s in a small, low-speed “picofuge”, rotated 180°, and spun again for ∼30 s to pack the beads on the bottom of the tube. After withdrawing a sample of the supernatant for SDS-PAGE analysis, the beads were washed twice with lysis buffer, aspirated thoroughly, and resuspended in 50 µl of 1x SDS loading buffer without dye. 10 µl each of insoluble pellet, lysate, bead supernatant, and bead-bound fractions were resolved on a 10% Bis-Tris SDS-PAGE gel and imaged as described above.

For size-exclusion chromatography (Figure S4B), lysates were separated on a Superdex 200 10/300 GL analytical column in 1-ml fractions, which were analyzed as above by SDS-PAGE.

### RNase A bead-loading

For RNase A treatment experiments (Figures 5C and S5G), cells that had been labeled overnight with JF dyes were washed twice with 1x PBS and destained for 30 min in phenol red-free DMEM. The medium was thoroughly aspirated and replaced with 20 µl of 1x PBS containing 5 mg/ml of RNase A (Qiagen) and 12 mM of purified H6-mNeonGreen-NLS protein as a loading control. The same solution without RNase A was used for the negative control condition. The cells were sprinkled with a thin layer of ≤ 106 µm glass beads (Sigma G4649) as described elsewhere,^81^ and the dish was tapped firmly ten times on the benchtop. Beads were rinsed away by washing several times with 1x PBS, and the cells were allowed to recover for 30 min in phenol red-free DMEM. The medium was changed once more before imaging.

To produce H6-mNeonGreen-NLS protein, BL21 (DE3) *E. coli* cells transformed with an expression plasmid were grown in 1 liter of LB with 100 µg/ml carbenicillin to an OD600 of 0.5 and induced with 1 mM isopropyl β-D-thiogalactopyranoside (IPTG) for 6 h at 37°C. Cells were pelleted, flash-frozen in liquid nitrogen, and stored at −80°C. Cells were lysed by sonication in lysis buffer (20 mM Tris-HCl, pH 8, 1 M NaCl, 30 mM imidazole, 1 mM DTT, 1 mM benzamidine, and one EDTA-free complete protease inhibitor cocktail tablet [Roche] per 50 ml). The lysate was clarified by centrifugation at 38,000 *g* in a fixed-angle rotor, and the supernatant was incubated with 0.5 ml of NiNTA agarose (Qiagen) for several hours with rotation at 4°C. The slurry was poured into a disposable polypropylene column, and the resin was washed with lysis buffer followed by wash buffer (10 mM Tris-HCl, pH 7.5, 100 mM NaCl, 30 mM imidazole, 1 mM DTT). Protein was eluted with elution buffer (10 mM Tris-HCl, pH 7.5, 100 mM NaCl, 250 mM imidazole, 1 mM DTT). Eluted protein was diluted with 2.5 volumes of SP buffer (10 mM Tris, pH 7.5, 1 mM DTT) to reduce its ionic strength, loaded onto a HiTrap SP column, and eluted with a linear gradient of 100 mM to 1 M NaCl in SP buffer. Elution was monitored by measuring absorbance at 260, 280, and 506 nm. Peak fractions containing protein were pooled and dialyzed against 1x PBS with 1 mM DTT. Single-use aliquots were flash-frozen and stored at −80°C.

### Tat expression experiment

To test the effect of Tat on HEXIM1:CycT1 interaction (Figure 6A), HEXIM1-Halo + SNAPf-CycT1 U2OS cells (clone SC-B12) were trypsinized and counted, and 1.2 million cells were electroporated with 1 µg of either mNeonGreen-NLS (pTG1033) or Tat-mNeonGreen (pTG1035) expression plasmid, using the Amaxa Nucleofector system with Lonza Kit V solutions. The cells were divided between two glass-bottom dishes (MatTek P35G-1.5-20-C) and stained overnight with 50 nM of JFX549 HaloTag ligand and 5 nM of JFX650 SNAP ligand in phenol red-free DMEM. PAPA was measured using the automated protocol described above. For each field of view, an additional snapshot was taken with 488 nm excitation to measure mNeonGreen signal.

## Author contributions

All the authors contributed to conceptualization of the project. G.M.D. designed the new plasmids used in this work, and G.M.D. and B.W. did molecular cloning. C.D.D., G.M.D., B.W., and T.G.W.G. generated and validated U2OS cell lines. T.G.W.G. performed the experiments shown in the figures and wrote the manuscript. B.W. assisted with the experiment in Figure 6A. T.G.W.G., R.T., C.D.D., and X.D. edited the manuscript.

---

**Table S1:** fSMT sample sizes.

| Figure | Condition | Number of biological replicates | # movies | # single-molecule trajectories |
| --- | --- | --- | --- | --- |
| 2A, S1C | HEXIM1-SNAPf, clone S-A6 | 4 | 156 | 346205 |
| 2B, S1B | SNAPf-CycT1, clone G4-9 | 7 | 239 | 341406 |
| 4A and S5A,C | HEXIM1-SNAPf, clone S-A6 + NVP-2 | 2 | 242 (Figure S5A,C) | 559970 |
|  | -60 to -40 min |  | 22 | 49142 |
|  | -40 to -20 min |  | 23 | 51277 |
|  | -20 to 0 min |  | 21 | 48142 |
|  | 0 to 20 min |  | 21 | 51663 |
|  | 20 to 40 min |  | 20 | 49755 |
|  | 40 to 60 min |  | 21 | 44439 |
| 4B and S5B,D | SNAPf-CycT1, clone G4-9 + NVP-2 | 7 | 374 (Figure S5B,D) | 719698 |
|  | -60 to -40 min |  | 23 | 48425 |
|  | -40 to -20 min |  | 66 | 136379 |
|  | -20 to 0 min |  | 66 | 113792 |
|  | 0 to 20 min |  | 67 | 129901 |
|  | 20 to 40 min |  | 60 | 101339 |
|  | 40 to 60 min |  | 28 | 66812 |
| S1B | Halo-CycT1, clone G1 | 3 | 528 | 159932 |
| S1B | Halo-CycT1, clone F8 | 2 | 122 | 33435 |
| S1B | <u>Halo-CycT1</u> + HEXIM1-SNAPf, clone HC-C6 | 2 | 431 | 82327 |
| S1B | SNAPf-CycT1, clone H4-10 | 3 | 305 | 467522 |
| S1B | <u>SNAPf-CycT1</u> + HEXIM1-Halo, clone SC-B12 | 3 | 254 | 389534 |
| S1C | HEXIM1-Halo, clone H-G3 | 3 | 133 | 542595 |
| S1C | HEXIM1-Halo, clone H-G7 | 2 | 55 | 198588 |
| S1C | HEXIM1-Halo, clone H-D3 | 3 | 478 | 855992 |
| S1C | HEXIM1-SNAPf, clone S-D12 | 2 | 61 | 239771 |
| S3C | LARP7-Halo, clone HL-FF2 | 2 | 366 | 37434 |
| S3D | SNAPf-3xNLS | 1 | 91 | 34855 |

Notes:

**Table S2:** PAPA-fSMT sample sizes.

| Figure | Condition | # movies |  | # single-molecule trajectories |
| --- | --- | --- | --- | --- |
| 2C | Halo-CycT1 → HEXIM1-SNAPf | 612 | PAPA | 67810 |
|  |  |  | DR | 73312 |
| 2D | HEXIM1-Halo → SNAPf-CycT1 | 326 | PAPA | 71192 |
|  |  |  | DR | 27855 |
| 2E | LARP7-Halo → HEXIM1-SNAPf | 273 | PAPA | 52375 |
|  |  |  | DR | 46139 |
| 2F | LARP7-Halo → SNAPf-CycT1 | 180 | PAPA | 30354 |
|  |  |  | DR | 37705 |
| 5A | LARP7-Halo → hnRNP R (transgene) | 248 | PAPA | 36078 |
|  |  |  | DR | 30884 |
| 5A | LARP7-Halo → hnRNP Q-SNAPf | 206 | PAPA | 26872 |
|  |  |  | DR | 21103 |
| 5A | LARP7-Halo → hnRNP A1 | 115 | PAPA | 13352 |
|  |  |  | DR | 23446 |
| 5A | LARP7-Halo → hnRNP H1 | 258 | PAPA | 7414 |
|  |  |  | DR | 19275 |
| 5A | LARP7-Halo → hnRNP C2 | 164 | PAPA | 3809 |
|  |  |  | DR | 10659 |
| 5C | LARP7-Halo → hnRNP R (endogenous) | 143 | PAPA | 5188 |
|  |  |  | DR | 8115 |
| 5C | LARP7-Halo → hnRNP R (endogenous) + RNase A | 126 | PAPA | 4408 |
|  |  |  | DR | 15191 |
| 6B | Halo-BRD4 → SNAPf-CycT1 | 370 | PAPA | 18457 |
|  |  |  | DR | 34527 |
| S5F | LARP7-Halo → SNAPf-hnRNP Q | 424 | PAPA | 11563 |
|  |  |  | DR | 23289 |

**Table S3:** List of plasmids. Plasmid maps can be found at https://github.com/tgwgraham/ptefb_papa_plasmids.

| Stock number | Description | Source |
| --- | --- | --- |
| pTG755 | EF1αprom-SNAPf-3xNLS PB Puro | Ref. <sup>82</sup> |
| pTG888 | CMV-Renilla | Promega |
| pTG988 | EF1αprom-H2B-SNAPf-3xNLS PB Puro | Ref. <sup>82</sup> |
| pTG1033 | EF1αprom-mNeonGreen-NLS PB Puro | This work |
| pTG1035 | EF1αprom-HIV Tat-mNeonGreen PB Puro | This work |
| pTG1036 | EF1αprom-HIV Tat ZFnull-mNeonGreen PB Puro (all cysteines mutated to glycine) | This work |
| pTG1043 | pEF-Myc-Tat | Koh Fujinaga, UCSF |
| pTG1044 | HIV LTR luciferase | Koh Fujinaga, UCSF |
| pTG1148 | U6-hCCNT1 gRNA #2N CBh-Cas9 PGK-Venus | This work |
| pTG1149 | U6-hCCNT1 gRNA #3N CBh-Cas9 PGK-Venus | This work |
| pTG1150 | U6-hCCNT1 gRNA #4N CBh-Cas9 PGK-Venus | This work |
| pTG1151 | pHR-V5-SNAPf-CCNT1 #2N | This work |
| pTG1152 | pHR-V5-SNAPf-CCNT1 #3N | This work |
| pTG1153 | pHR-V5-SNAPf-CCNT1 #4N | This work |
| pTG1154 | U6-hHEXIM1 gRNA #1C CBh-Cas9 PGK-Venus | This work |
| pTG1155 | U6-hHEXIM1 gRNA #2C CBh-Cas9 PGK-Venus | This work |
| pTG1156 | U6-hHEXIM1 gRNA #4C CBh-Cas9 PGK-Venus | This work |
| pTG1157 | pHR-HEXIM1-SNAPf-V5 #1C | This work |
| pTG1158 | pHR-HEXIM1-SNAPf-V5 #2C | This work |
| pTG1159 | pHR-HEXIM1-SNAPf-V5 #4C | This work |
| pTG1201 | pHR_gRNA#2N_3XF-Halo-GDGAGLIN-hCCNT1 | This work |
| pTG1202 | pHR_gRNA#3N_3XF-Halo-GDGAGLIN-hCCNT1 | This work |
| pTG1203 | pHR_3XF-Halo-GDGAGLIN-hCCNT1_WT=4N | This work |
| pTG1204 | pHR_gRNA#1-2C_hLARP7-GDGAGLIN-Halo-3XF | This work |
| pTG1205 | pHR_gRNA#3C_hLARP7-GDGAGLIN-Halo-3XF | This work |
| pTG1206 | pU6_hLARP7_gRNA#1C(T5A)_CBh_Cas9_PGK_Venus | This work |
| pTG1207 | pU6_hLARP7_gRNA#2C(T5A)_CBh_Cas9_PGK_Venus | This work |
| pTG1208 | pU6_hLARP7_gRNA#3C_CBh_Cas9_PGK_Venus | This work |
| pTG1209 | pHR_gRNA#1C_hHEXIM1-GDGAGLIN-Halo-3XF | This work |
| pTG1210 | pHR_gRNA#2C_hHEXIM1-GDGAGLIN-Halo-3XF_GA | This work |
| pTG1211 | pHR_gRNA#4C_hHEXIM1-GDGAGLIN-Halo-3XF | This work |
| pTG1212 | pHR_gRNA#1N_3XF-Halo-GDGAGLIN-hBRD4_WT | This work |
| pTG1213 | pHR_gRNA#2N_3XF-Halo-GDGAGLIN-hBRD4 | This work |
| pTG1214 | pHR_gRNA#4N_3XF-Halo-GDGAGLIN-hBRD4 | This work |
| pTG1215 | pU6_hBRD4_gRNA#1N_CBh_Cas9_PGK_Venus | This work |
| pTG1216 | pU6_hBRD4_gRNA#2N_CBh_Cas9_PGK_Venus | This work |
| pTG1217 | pU6_hBRD4_gRNA#4N_CBh_Cas9_PGK_Venus | This work |

