## Supplemental figures for "Single-molecule live imaging of subunit interactions and exchange within cellular regulatory complexes"

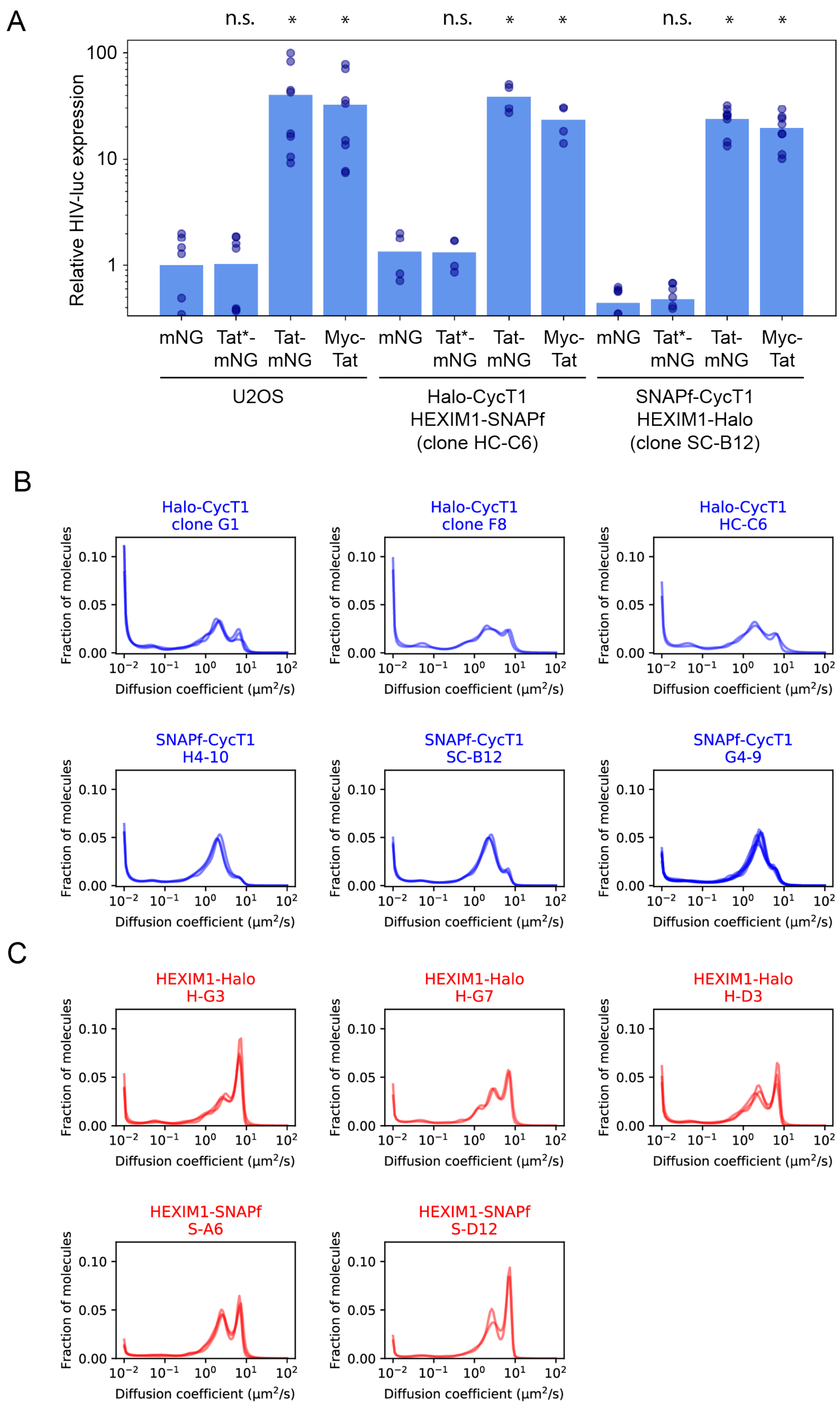

**Figure S1**

A) Tat induces the HIV LTR promoter in cell lines with Halo/SNAPf-tagged endogenous CycT1 and HEXIM1. HIV LTR-luciferase reporter signal in U2OS cells and Halo/SNAPf-tagged U2OS cell lines co-transfected with the indicated plasmids. mNG: mNeonGreen fluorescent protein (negative control). Tat*-mNG: zinc finger mutant Tat, with all cysteines mutated to alanine, fused to mNeonGreen. Tat-mNG and Myc-Tat: HIV Tat activator fused to mNeonGreen or Myc epitope tag, respectively. Relative HIV-luc expression is defined as the ratio of HIV-luciferase to CMV-Renilla luciferase internal control, normalized to the mean of the U2OS mNeonGreen negative control condition. A Bonferroni-corrected, 2-tailed t-test in log space was used to compare each condition to the reporter-only condition for that cell line. *, $p < 10^{-4}$. n.s., $p > 0.05$.

B) State array analysis of fast single-molecule tracking (fSMT) data for multiple endogenously tagged CycT1 clonal lines. While Halo- and SNAPf-tagged CycT1 clonal lines showed peaks at similar diffusion coefficients, the proportion of molecules in the 2-3 $\mu m^2/s$ peak was higher for the three SNAPf-CycT1 clonal lines than for the three Halo-CycT1 clonal lines, suggesting that fusion of CycT1 to Halo and/or SNAPf may influence its distribution between different complexes. Data for SNAPf-CycT1 clone G4-9 are reproduced here from Figure 2B.

C) State array analysis of fSMT data for multiple endogenously tagged HEXIM1 clonal cell lines. Data for HEXIM-SNAPf clone S-A6 are reproduced here from Figure 2A.

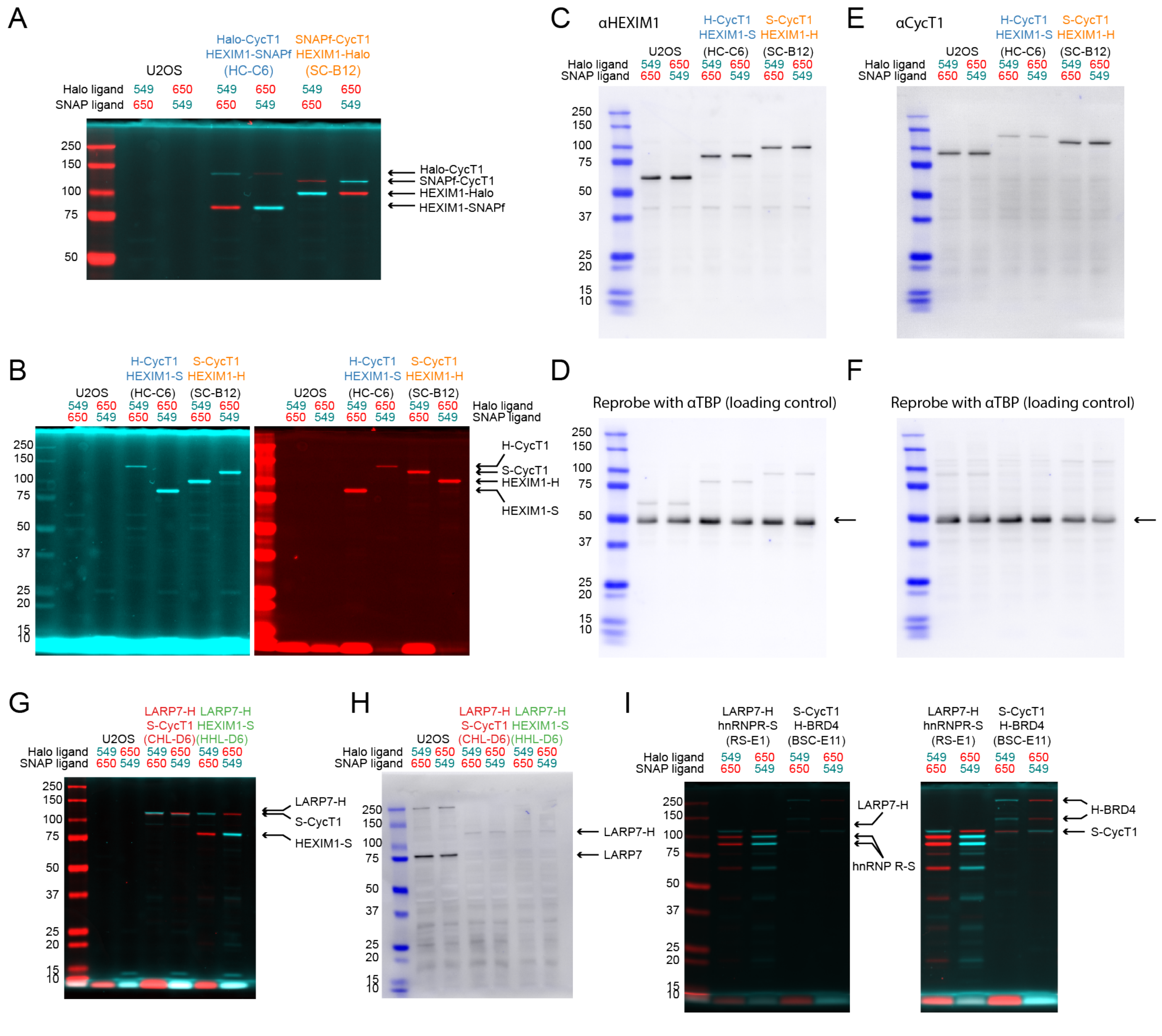


**Figure S2 - related to Figures 2 and 3**

Validation of double-labeled cell lines by SDS-PAGE and western blotting.

A) SDS-PAGE gel of lysates prepared from the indicated cell lines stained with JF(X)549/JFX650 Halo or SNAPf ligands, as indicated (also see Methods). Shown is an overlay of the JF(X)549 (cyan) and JFX650 (red) channels. Molecular weight in kilodaltons is indicated on the left. The identities of labeled proteins are indicated on the right. Ladder bands are saturated in the JFX650 channel.

B) Full-size single-channel images of the gel in (A) with increased contrast, showing the absence of lower-molecular weight degradation products. The display of the brightest bands is saturated.

C) Western blot of the same lysates from (A,B) with an anti-HEXIM1 primary antibody. HRP chemiluminescence (black) is overlaid with far-red fluorescence of the ladder (blue).

E) Western blot of the same lysates from (A,B) with an anti-CycT1 primary antibody.

D,F) Western blots from (C) and (E), respectively, reprobed with an anti-TATA binding protein (TBP) antibody as a loading control (arrow).

G) SDS-PAGE gel of lysates from LARP7-Halo + SNAPf-CycT1 and LARP7-Halo + HEXIM1-SNAPf cells stained with JF(X)549/JFX650 Halo or SNAP ligands. Cyan: JF(X)549. Red: JFX650. Molecular weight in kilodaltons is indicated on the left, and band assignments are shown on the right.

H) Anti-LARP7 western blot of lysates from (G). Successful Halo-tagging of LARP7 is indicated by the disappearance of the wild-type LARP7 band and the appearance of a higher-molecular weight band. We do not know whether weaker signal from the Halo-LARP7 band compared to untagged LARP7 reflects a reduced expression level, poorer transfer to the nitrocellulose membrane, or poorer detection efficiency by the antibody.

I) SDS-PAGE gel of fluorescently labeled proteins in lysates of LARP7-Halo + hnRNP R-SNAPf (clone RS-E1) and SNAPf-CycT1 + Halo-BRD4 (clone BSC-E11) cell lines. The same image is shown on the right with higher contrast to make faint bands visible. The sizes of the two major bands of hnRNP R and BRD4 are consistent with previously annotated short and long isoforms of these two proteins. Fainter, lower molecular weight bands observed for hnRNP R may be degradation products.

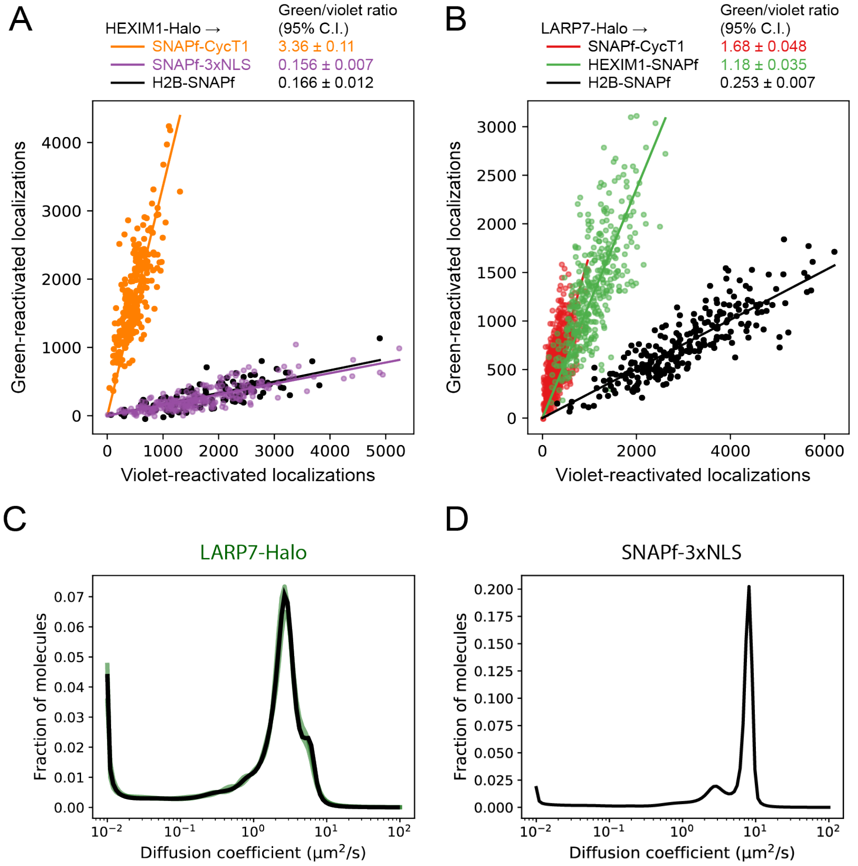


**Figure S3 - related to Figures 2 and 3**

A-B) PAPA signal between labeled CycT1, HEXIM1, and LARP7 is greater than nonspecific background reactivation. Scatterplots of total green-reactivated (PAPA) and violet-reactivated (DR) localizations for different sender-receiver pairs. The number of reactivated localizations is defined as the difference in total number of localizations before and after each reactivation pulse. The green/violet (GV) ratio, defined as the ratio of green-reactivated to violet-reactivated localizations, provides a normalized interaction signal.

A) JFX549 sender-labeled HEXIM1-Halo with JFX650 receiver-labeled SNAPf-CycT1 (orange), nuclear SNAPf (SNAPf-3xNLS; lavender) and SNAPf-tagged histone H2B (black). SNAPf-3xNLS and H2B-SNAPf served as non-interacting negative controls.

B) JFX549 sender-labeled LARP7-Halo with JFX650 receiver-labeled SNAPf-CycT1 (red), HEXIM1-SNAPf (green), or non-interacting H2B-SNAPf negative control (black).

C) fSMT of LARP7. Green lines: individual experimental replicates. Solid black line: all data combined.

D) fSMT of SNAPf tag only with a 3x SV40 nuclear localization sequence (SNAPf-3xNLS).

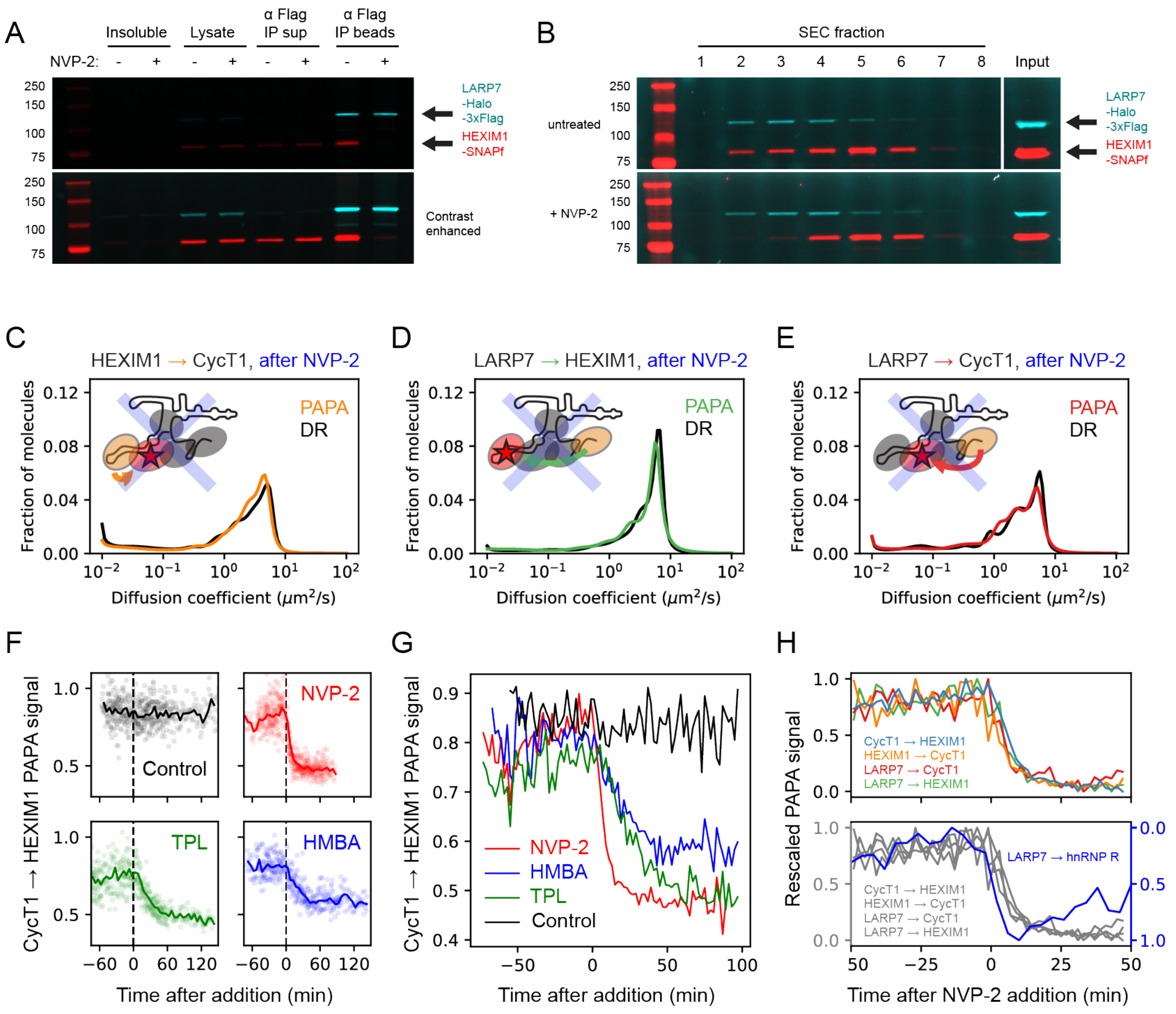


**Figure S4 - related to Figures 3 and 5.**

A-B) Endpoint assays demonstrate dissociation of labeled HEXIM1 from LARP7 in cell lysates. HEXIM1-SNAPf + LARP7-Halo-3xFlag double knock-in cells were labeled with JFX549 Halo ligand and JFX650 SNAPf ligand and either treated with NVP-2 for 30 min or left untreated. Cell lysates were prepared and subjected to either A) immunoprecipitation with an anti-Flag antibody or B) size-exclusion chromatography (SEC). Fractions were separated by SDS-PAGE and imaged on a fluorescence imager. NVP-2 treatment of cells eliminated co-immunoprecipitation of HEXIM1 with LARP7 (A, last lane) and shifted HEXIM1 toward smaller-molecular weight SEC fractions (B; compare HEXIM1-SNAPf in fractions 2-3 between +/- NVP2 conditions). Note that image contrast is enhanced in (B) to make visible the difference in HEXIM1-SNAPf levels in fractions 2 and 3.

C-E) Diffusion profiles of PAPA and DR trajectories in cells treated with NVP-2 for at least 20 min. In contrast to untreated cells (Figure 2D-F), there is no appreciable shift between the diffusion profiles of DR and PAPA trajectories, consistent with the interpretation that residual PAPA signal (Figure 3) is nonspecific.

F-G) PAPA detects dissociation of CycT1 from HEXIM1 in response to multiple drugs. PAPA was measured between JFX549-Halo-CycT1 and HEXIM1-SNAPf-JFX650 as in Figure 3A, and different compounds were applied to cells during the imaging time course. Triptolide (TPL) blocks transcription initiation by inhibiting the TFIIH helicase, while N,N'-hexamethylene bis(acetamide) (HMBA) activates Akt signaling.[1] F) Time courses of CycT1 → HEXIM1 PAPA signal (GV

ratio) upon addition of different drugs. Each point represents a single cell, and solid lines show averages over 2-min time bins. NVP-2 data are reproduced from Figure 3A. G) Overlay of binned averages from (F).
H) Top panel: Overlay of average traces from NVP-2 treatment experiment in Figure 3, rescaled to minimum and maximum PAPA signal. Bottom panel: Traces from the top panel (gray) overlaid with inverted and rescaled LARP7 → hnRNP R from Figure 5D.

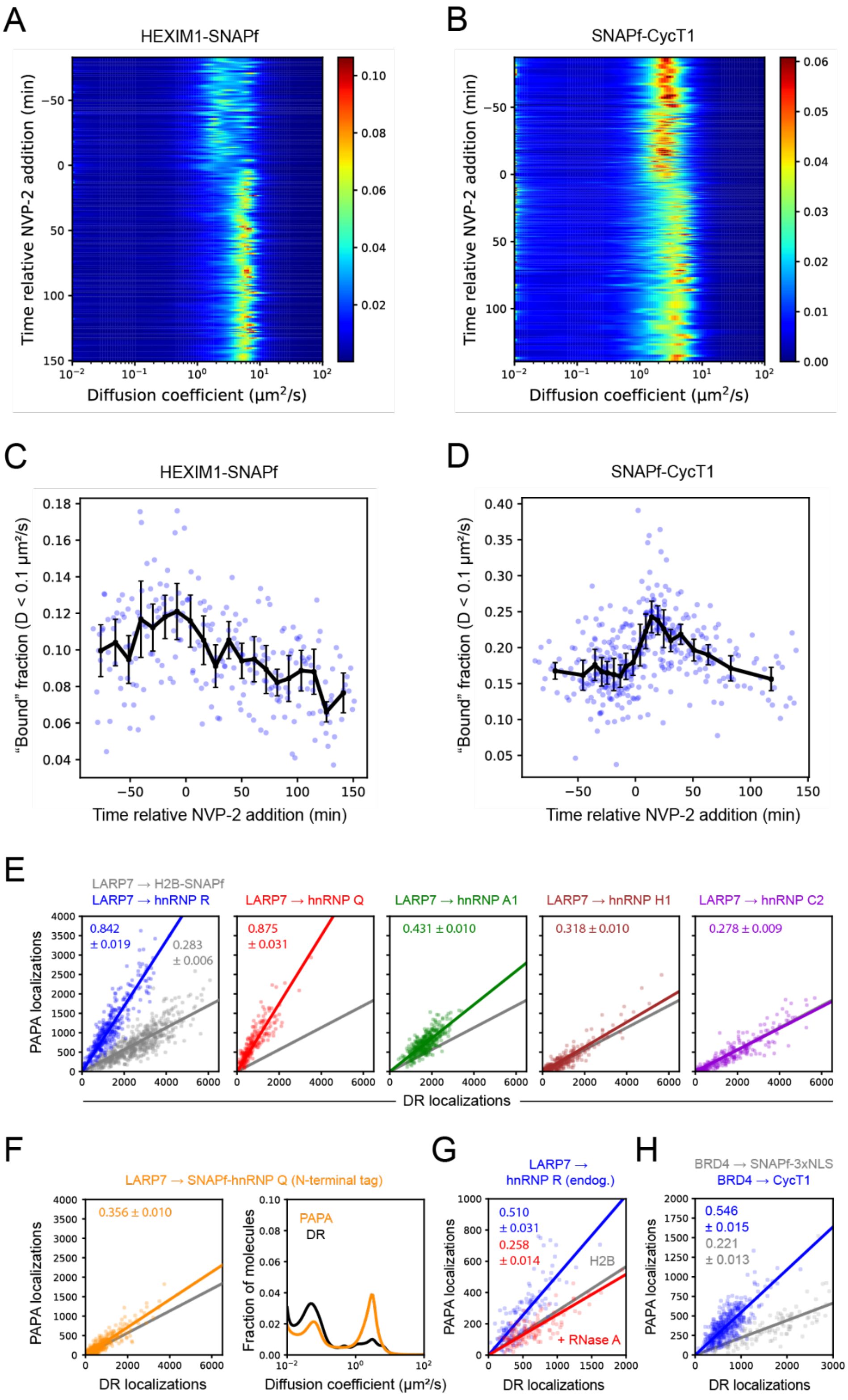

**Figure S5 - related to Figures 4, 5, and 6**

A-B) Diffusion profiles of HEXIM1-SNAPf (A) and SNAPf-CycT1 (B) molecules within individual cells as a function of time before or after addition of the Cdk9 inhibitor NVP-2. Each row corresponds to a distinct cell, and the color map represents the fraction of molecules at each diffusion coefficient. A shift in the diffusion profile of both proteins is observed upon inhibitor addition.

C-D) "Bound" fraction ($D < 0.1\ \mu m^2/s$) of HEXIM1-SNAPf (C) and SNAPf-CycT1 (D) molecules as a function of time before or after NVP-2 addition.

E) Related to Figure 5A-B. Increase in the number of localizations induced by direct reactivation (DR) of different SNAPf-tagged hnRNPs (horizontal axis) and PAPA from LARP7-Halo (vertical axis). Color points represent individual cells expressing SNAPf-tagged hnRNPs, while gray points in the first panel represent individual cells expressing H2B-SNAPf as a negative control. Solid lines show linear fits constrained to pass through the origin. The H2B-SNAPf control line is reproduced in every panel for reference. A slope greater than that of H2B-SNAPf indicates association above background.

F) PAPA vs. DR localization plot (as in E) and PAPA vs. DR diffusion profiles (as in Figure 5A) for N-terminally SNAPf-tagged hnRNP Q. The LARP7 → H2B-SNAPf negative control (gray line) is reproduced from panel (E).

G) PAPA vs. DR localization plot for the RNase treatment experiment shown in Figure 5C. The LARP7 → hnRNP R PAPA signal dropped to the background level of LARP7 → H2B (reproduced from panel (E)) upon treatment with RNase A, indicating the loss of specific interaction.

H) PAPA vs. DR localization plot for the BRD4 → CycT1 experiment in Figure 6B. BRD4 → SNAPf-3xNLS is included as a negative control.

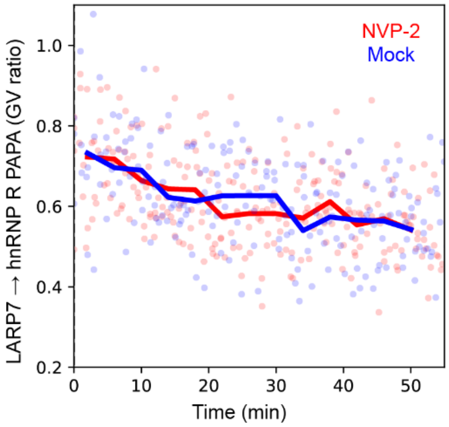


**Figure S6 – related to Figure 5D**

LARP7 → hnRNP R PAPA signal as a function of time after the start of imaging. For unknown reasons, we observed a downward drift in the PAPA signal at the beginning of each time course. NVP-2 or DMEM only as a negative control were added after the GV ratio had reached a plateau (i.e., after the time range displayed in this plot), and the first 30 time points of each time course are excluded from the plot in Figure 5D.

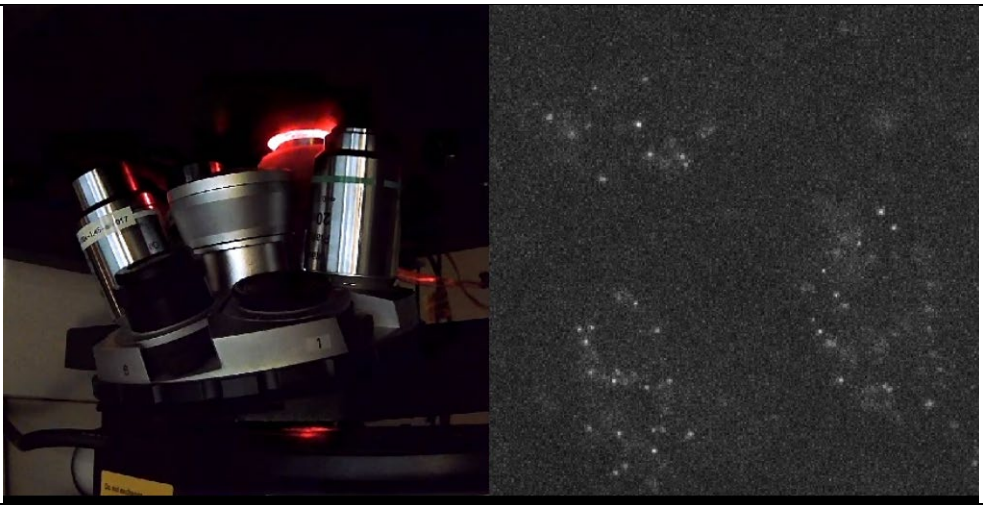

**Movie S1 – related to Figure 1**
PAPA-fSMT illumination sequence. Left panel shows a view underneath the microscope with laser light exiting the objective lens. Right panel shows single-molecule images acquired at the same time by the EMCCD camera. Each movie involves multiple cycles of PAPA (green light stimulation) and DR (violet light stimulation) separated by bleaching periods (continuous red illumination). Molecules are imaged with stroboscopic red illumination, and EMCCD acquisition is paused during PAPA, DR, and bleaching pulses. The 48 x 48 µm field of view shown here, containing three cell nuclei, is larger than the typical single-nucleus field of view that we used for fast tracking. The video is shown at 1x speed.
